# Global genomic epidemiology of multi-carbapenemase Enterobacterales

**DOI:** 10.64898/2026.10.07.757280

**Authors:** Nicole Lerminiaux, Dorottya Nagy, Sam Lipworth, the NEKSUS Consortium, Liam P. Shaw, Laura Mataseje, Katie L. Hopkins, Kevin Chau, Nicole Stoesser

## Abstract

Enterobacterales pathogens that encode multiple carbapenemase genes are an emerging public health threat. The genomic and plasmid context of the multi-carbapenemase producers has not yet been explored at scale. We analysed 1501 publicly-available global genomes that encoded two or more genes from the five prevalent carbapenemase families (IMP, KPC, NDM, OXA-48-like, and VIM). We observed 241 distinct allelic combinations, with NDM/OXA-48-like (25.6%), multiple KPC (25.3%), KPC/NDM (18.8%), and multiple NDM (10.9%) being the most common groups. We found that many genomes were part of high-risk lineages (48.0%). Most genomes carried carbapenemase genes on two plasmids (59.2%) or one plasmid (23.5%), and most genomes encoded different carbapenemase families (56.2%) rather than copies of the same allele (36.2%). We observed multiple mechanisms of multi-carbapenemase evolution including plasmid co-integration, convergence with virulence, and generation of novel allele diversity through duplication. Multi-carbapenemase plasmids had more AMR genes that conferred resistance to more AMR classes compared to plasmids encoding a single carbapenemase. The proportion of cross-class combinations (KPC/NDM and NDM/OXA-48-like) increased significantly over time, with NDM-5/OXA-181 (3.3-fold) and NDM-4/OXA-181 (8.9-fold) combinations enriched. These findings provide valuable insights into the genomic features of multi-carbapenemase Enterobacterales to support the development of effective surveillance programmes.

## INTRODUCTION

Carbapenemase genes confer resistance to carbapenem antibiotics, which are last-resort treatment options for multi-drug-resistant Enterobacterales infections. Carbapenem-resistant Enterobacterales are listed as Critical on the 2024 World Health Organization Priority Pathogens List^1^, and their worldwide dissemination poses a major challenge to public health^2,3^. Carbapenemase genes are frequently carried on mobile genetic elements that facilitate their mobility^4,5^ and complicate clinical outbreak investigations^6^.

The major carbapenemase families (IMP, KPC, NDM, OXA-48-like, VIM) have varying hydrolytic profiles and inhibitor susceptibilities and therefore require different clinical treatment strategies^7^. However, the emergence of isolates encoding multiple carbapenemase genes is complicating treatment^8–15^. Multi-carbapenemase isolates have broader antimicrobial resistance (AMR) profiles than single carbapenemase producers^11,13^ and harbouring multiple copies of the same carbapenemase gene can result in higher minimum inhibitory concentrations to antibiotics such as ceftazidime and ceftazidime-avibactam^16^. Several recent studies have described lineages and temporal trends in multi-carbapenemase gram-negative bacteria^13^ and in *Klebsiella* encoding KPC/NDM specifically^14,17^, yet the genomic and plasmid context of multi-carbapenemase producers has not yet been explored at scale. In addition, it is unclear if multiple carbapenemase genes are found in the same genome solely due to selective pressures (co-selection) or if they are physically linked on the same genetic element in close proximity (co-mobilisation/genetic linkage), enabling them to be co-transferred^18,19^.

To characterize the genomic features and plasmids of multi-carbapenemase isolates, we analysed publicly available complete genomes carrying two or more genes from the most prevalent carbapenemase families (IMP, KPC, NDM, OXA-48-like, VIM). Using complete genomes allowed us to investigate the genomic location and context of carbapenemase genes across diverse allele combinations and lineages. This study provides insight into the global genomic epidemiology of multi-carbapenemase isolates to inform effective surveillance and the development of future treatment strategies.

## RESULTS

### A global dataset of multi-carbapenemase genomes

We identified 1501 multi-carbapenemase genomes isolated from 2008 – 2025 (**Supplementary Table 1**). We defined “multi-carbapenemase” as two or more copies of the same or different carbapenemase genes. Of the Enterobacterales genomes in NCBI GenBank with ≤ 20 contigs (n=46531), 22.9% (10676/46531) encoded at least one carbapenemase gene and 14.1 % (1501/10676) of these encoded multiple carbapenemase genes, resulting in multi-carbapenemase genomes comprising 3.2 % (1501/46531) of the available genomes (**Supplementary Figure 1**).

Most multi-carbapenemase genomes were from Asia (50.1%, 752/1501), Europe (21.2%, 318/1501), or North America (20.7%, 311/1501) (**Supplementary Figure 2**). Africa (1.5%, 23/1501), Oceania (1.3%, 20/1501), and South America (2.7%, 41/1501) genomes were not well represented in this dataset. Multi-carbapenemase genomes were found in 12 genera and the most prevalent species were within the *Klebsiella pneumoniae* species complex (58.6%, 879/1501), followed by *Escherichia coli* (13.9%, 209/1501). Multi-carbapenemase *Kluyvera, Leclercia, Morganella, Proteus, Providencia, Salmonella,* and *Serratia* were represented by fewer than 30 genomes each.

The genomes encoded a combined total of 3323 carbapenemase genes (**Supplementary Table 2, Supplementary Table 3**), with most genomes carrying two (85.9%, 1290/1501) or three (10.5%, 158/1501) carbapenemase genes. NDM/OXA-48-like (25.6%, 384/1501), multiple KPC (25.3%, 380/1501), KPC/NDM (18.8%, 282/1501), and multiple NDM (10.9%, 163/1501) were the most common carbapenemase groups (**Figure 1**). There were 241 distinct allelic combinations, with the most common combinations being KPC-2/NDM-1 (10.8%, 162/1501), KPC-2/KPC-2 (10.3%, 155/1501), KPC-3/KPC-3 (5.7%, 86/1501), NDM-1/OXA-48 (5.2%, 78/1501), and NDM-5/OXA-181 (5.0%, 75/1501) (**Supplementary Figure 3, Supplementary Figure 4**). IMP and VIM carbapenemases were relatively rare (**Supplementary materials**). The genomes also encoded other AMR genes that conferred resistance to a median of nine antimicrobial classes (range: 1 – 15), with *sul1* (2292 copies, 77.1% (1158/1501) of genomes) and *bla*_TEM-1_ (1292 copies, 65.3% (980/1501) of genomes) found most frequently.

**Figure 1.**
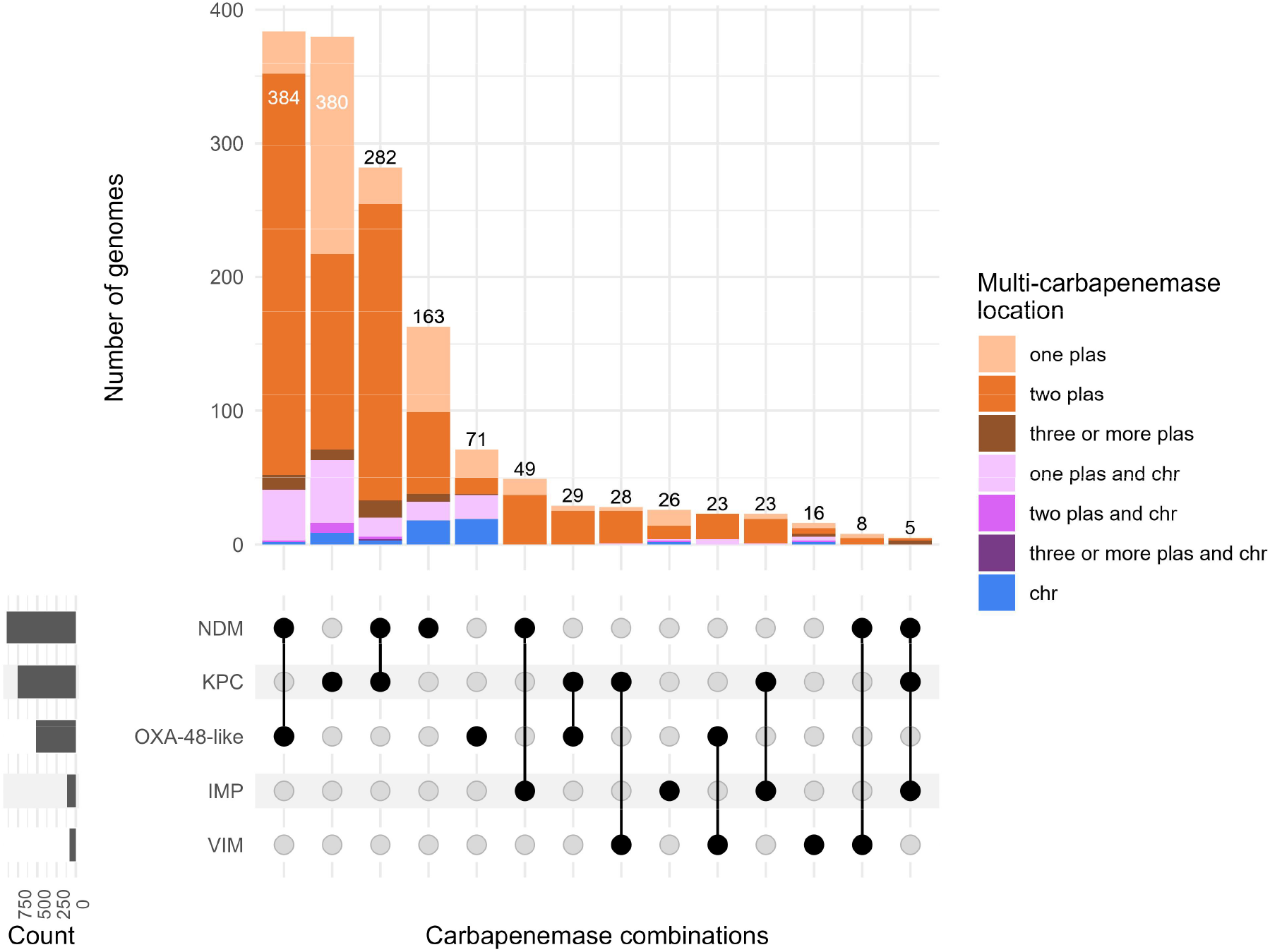
**Most common carbapenemase groups coloured by genomic location**. “plas” = plasmid, “chr” = chromosome. Carbapenemase combinations in fewer than 5 genomes are not shown (n = 6 combinations representing 14 genomes; IMP/OXA-48-like n=4, IMP/VIM n=3, KPC/NDM/OXA-48-like n=3, KPC/NDM/VIM n=2, KPC/OXA-48-like/VIM n=1, NDM/OXA-48-like/VIM n=1).

### Diverse multi-carbapenemase gene carriage among high-risk species lineages suggests multiple acquisitions rather than clonal spread

High-risk lineages comprised 48.0% (721/1501) of genomes in this dataset, and they predominantly carried multiple KPC (33.0%, 238/721), NDM/OXA-48-like (27.9%, 201/721), or KPC/NDM (20.0%, 144/721) (**Supplementary Table 1**). We observed diverse carbapenemase gene combinations within most high-risk clones (**Figure 2**), which suggests there have been multiple independent acquisitions of multi-carbapenemases in each lineage as opposed to clonal dissemination. The exception was *E. hormaechei* ST79, in which NDM-5/OXA-48-carrying isolates were linked to a clonal outbreak in Denmark^20^.

**Figure 2.**
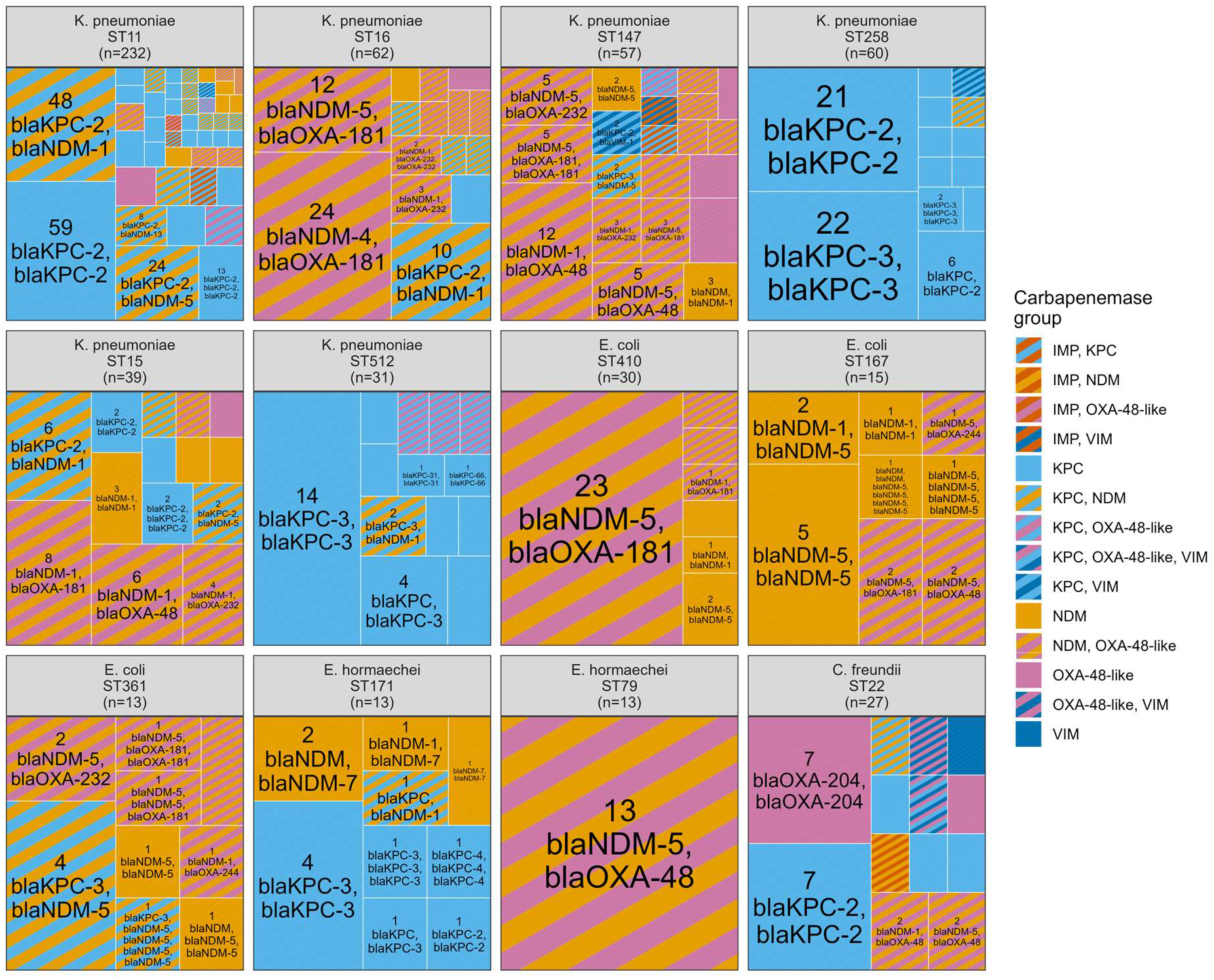
Most common carbapenemase allelic combinations found across high-risk lineages. Size of squares are scaled to the proportion of each category.

*K. pneumoniae* sequence type (ST) 11 was the most frequent lineage among the *K. pneumoniae* complex (26.4%, 232/879) and had the highest number of carbapenemase gene combinations (n=46). The most common combinations were KPC-2/KPC-2 (12.5%, 59/232) and KPC-2/NDM-1 (20.7%, 48/232). In contrast, single-locus variants ST258 (6.8%, 60/879) and ST512 (3.5%, 31/879) harboured predominantly multiple copies of KPC. Other high-risk *K. pneumoniae* lineages (ST147: 6.5%, 57/879; ST15: 4.4%, 39/879; ST16: 7.1%, 62/879) most often carried NDM/OXA-48-like. In other genera, *E. coli* ST410 (14.4%, 30/209), ST167 (7.2%, 15/209), and ST361 (6.2%, 13/209) were found most frequently among *E. coli* (**Figure 2**). ST410 genomes had NDM-5/OXA-181 as the dominant genotype (77%, 23/30), whereas ST167 genomes typically had multiple NDM copies (67%, 10/15) and ST361 genomes carried KPC-3/NDM-5 most frequently (31%, 4/13). *E. hormaechei* ST171 (11.2%, 13/116) primarily carried multiple KPC copies (46%, 6/13). *C. freundii* ST22 (26.2%, 27/103) was the most common *Citrobacter* lineage and it had the highest diversity of carbapenemase gene combinations outside the *Klebsiella* lineages (n=13).

Most carbapenemase gene combinations were predominantly found in *Klebsiella* due to its dominance in the dataset (**Supplementary Figure 5**); however, several combinations (NDM-5/OXA-181 and NDM-5/NDM-5) were significantly overrepresented in *E. coli* (NDM-5/OXA-181: 54/75, 72.0%, standardized residual (representing the magnitude of departure from the expected frequency)=14.9, BH-adjusted p<0.001; NDM-5/NDM-5: 19/33, 57.6%, standardized residual=7.3, BH-adjusted p<0.001; **Supplementary Table 4**). Combinations that were significantly overrepresented in *Klebsiella* included NDM-1/OXA-232 (39/39, 100%, standardized residual=4.8, BH-adjusted p<0.001), NDM-5/OXA-232 (38/40, 95.0%, standardized residual=4.2, BH-adjusted p<0.001), and KPC-2/NDM-5 (37/39, 94.8%, standardized residual=4.1, BH-adjusted p<0.001), whereas NDM-5/OXA-181 (20/75, 26.7%, standardized residual=-6.8, BH-adjusted p<0.001) and NDM-5/NDM-5 (11/33, 33.3%, standardized residual=-3.7, BH-adjusted p=0.004) were significantly underrepresented in *Klebsiella*.

### Different carbapenemase families are found on separate plasmids and multiple copies of the same carbapenemase gene are found on the same plasmid

We investigated the genomic location of the carbapenemase genes for all 1501 genomes (**Figure 1**). Most genomes carried carbapenemase genes on two plasmids (59.2%, 889/1501), followed by a single plasmid (23.5%, 353/1501). Some genomes had carbapenemase genes found on the chromosome in addition to one (9.5%, 142/1501) or two (0.8%, 12/1501) plasmids, and a minority had all carbapenemase genes chromosomally located (3.7%, 55/1501). Genomes with solely chromosomal carbapenemase genes typically encoded multiple OXA-48-like copies (35%, 19/55) or multiple NDM copies (33%, 18/55).

There was a significant association between the type of carbapenemase gene carried (same allele vs different allele vs different family) and genomic location (Fisher’s exact test, p<0.001). Genomes with different carbapenemase families (56.3%, 845/1501) were more likely to harbour them on two distinct plasmids (77.6%, 656/845), most consistent with multiple plasmid acquisition events, whereas genomes carrying copies of the same allele (36.2%, 544/1501) were more likely to harbour those genes on a single plasmid (40.1%, 218/544) or chromosome (8.8%, 48/544), most consistent with gene duplication/transposition events. It was rare to find genomes with different alleles for the same carbapenemase family (7.5%, 112/1501), and these were generally found on a single plasmid (41.1%, 46/112) or two plasmids (40.2%, 45/112).

We examined if there was an increase in multiple carbapenemase genes on single plasmids compared with other genomic locations over time, which would indicate an overall consolidation of carbapenemase genes onto individual plasmids. While there was a slight annual increase in the number of multiple carbapenemase genes on one plasmid, this was not statistically significant (Firth’s penalised binary logistic regression, OR=1.02, 95% CI [0.98, 1.06], p=0.28) (**Supplementary Table 5**).

We investigated if certain carbapenemases genes co-occurred more frequently than expected in a de-replicated subset of genomes to control for over-sampling of outbreak strains (see Methods, n=1217). Overall, duplicate copies of the same allele occurred more frequently than expected under the permutation null model (**Supplementary Table 6**); for example, multiple KPC-3 copies (8.0-fold enrichment, FDR-adjusted q=0.007) and multiple NDM-1 copies (4.3-fold enrichment, FDR-adjusted q=0.007) in the same genome were significantly enriched. Only two combinations of different families were significantly enriched, which were NDM-5/OXA-181 (3.3-fold enrichment, FDR-adjusted q=0.007) and NDM-4/OXA-181 (8.9-fold enrichment, FDR-adjusted q=0.007; 16/26 (61.5%) genomes occurred in *K. pneumoniae* ST16 from Asia and may represent a wider-scale outbreak).

A subset of genomes encoded three or more carbapenemase genes (14.1%, 211/1501), which encompassed 11 different genera (**Supplementary Figure 6**). Most genomes encoded either multiple copies of a single allele (46.0%, 97/211) or multiple copies of at least one allele alongside other genes (41.2%, 87/211). Few genomes encoded three or more distinct alleles without duplicates (12.8%, 27/211), and even fewer encoded three carbapenemase families (44%, 12/27; five IMP/KPC/NDM, three KPC/NDM/OXA-48-like, two KPC/NDM/VIM, one KPC/OXA-48-like/VIM, and one NDM/OXA-48-like/VIM), which were generally present on two (17%, 2/12) or three distinct plasmids (67%, 8/12). The exceptions were two closely related *Providencia hangzhouensis* ST222 isolates from North America that carried KPC-2, NDM-1, and VIM-24 on a single 190.9 kb IncC/rep_cluster_1254 plasmid. Eleven genomes had more than five carbapenemase genes (5.2%, 11/211), and in all but two cases, these were multiple copies of NDM (55%, 6/11) or KPC (18%, 2/11). No genomes encoded four or five carbapenemase families.

### Multi-carbapenemase plasmids encode more AMR genes and more AMR gene classes than single carbapenemase plasmids

We identified 6897 carbapenemase and non-carbapenemase plasmids from 1501 genomes. There were an additional 562 extra-chromosomal contigs identified as viruses, 13 of which (2.3%) contained carbapenemase genes; we excluded these from plasmid analysis but noted that 7/13 (54%) of these contigs had high identity to phage-plasmids SSU5 (n=5) and pKpn (n=2).

We assessed whether individual plasmids encoding two or more carbapenemase genes encoded additional AMR genes conveying resistance to other antibiotic classes compared with other plasmids (**Supplementary Figure 7, Supplementary Table 7**). To eliminate the influence of abundant small plasmids which typically do not encode AMR genes, we focused only on plasmids that encoded at least one AMR gene (**Table 1**; n=3857 plasmids overall). We observed statistically significant differences in AMR gene count and AMR gene classes between groups (Kruskal-Wallis for genes: χ^2^=103.8, p<0.001; for classes: χ^2^=79.0, p<0.001). Plasmids with multiple carbapenemase genes had a significantly higher number of AMR genes conveying resistance to other antibiotics (median: 6) compared with plasmids encoding only one (median: 3, Dunn’s test Z=8.6, BH-adjusted p<0.001) or zero carbapenemase genes (median: 4, Z=3.5, BH-adjusted p<0.001). Plasmids with multiple carbapenemase genes also had a higher number of AMR gene classes (median: 3) compared with single-carbapenemase plasmids (median: 2, Z=5.5, BH-adjusted p<0.001). In other words, on average a plasmid with an additional carbapenemase gene also has an additional three AMR genes conferring resistance to one additional antibiotic class. Plasmids with multiple carbapenemase genes were also significantly larger in size (median: 116.9 kb) and had a higher number of replicons (median: 2) than those with one carbapenemase gene (median: 83.6 kb, Z=8.1, BH-adjusted p<0.001 and median: 2, Z=6.7, BH-adjusted p<0.001, respectively). This supports the idea that most plasmids with two or more carbapenemase genes are likely hybrid co-integrates.

**Table 1.** Features of AMR-encoding plasmids (n=3857) with zero, one, or multiple carbapenemase genes from multi-carbapenemase genomes.

| Number of carbapenemase genes | Total plasmid count | Median number of AMR genes<sup>1</sup> [IQR<sup>2</sup>] | Median number of AMR gene classes<sup>1</sup> [IQR<sup>2</sup>] | Median number of replicons [IQR<sup>2</sup>] | Median length (kb) [IQR<sup>2</sup>] | Median number of TA systems [IQR<sup>2</sup>] |
| --- | --- | --- | --- | --- | --- | --- |
| None | 1410 | 4 [1-7] | 3 [1-5] | 1 [1-3] | 98.0 [45.7-176.2] | 1 [0-2] |
| One | 2025 | 3 [1-7] | 2 [1-5] | 2 [1-2] | 83.6 [51.5-139.7] | 0 [0-1] |
| Two or more | 422 | 6 [2-12] | 3 [1-7] | 2 [1-3] | 116.9 [77.7-181.2] | 0 [0-1] |

<sup>1</sup>Excluding carbapenemase genes; <sup>2</sup>IQR: interquartile range

To evaluate if carbapenemase plasmids were maintained by their hosts through addiction modules, we screened all plasmids for toxin-antitoxin (TA) systems. Approximately 61.4% (866/1410) of non-carbapenemase AMR-encoding plasmids and 50.9% (1246/2447) of carbapenemase plasmids encoded at least one TA system (n=3977, range: 1-9). Of the TA systems found on carbapenemase plasmids (53.4%, 2134/3977), the most common types were *higAB* (22.4%, 479/2134), *vagCD* (18.4%, 392/2134), *pemIK* (12.5%, 267/2134), and *relBE* (9.0%, 193/2134). When restricting the plasmid dataset to those that had at least one AMR gene, plasmids with zero carbapenemase genes had significantly more TA systems (median: 1) than plasmids with one (median: 0, Z=8.8, BH-adjusted p<0.001) or two or more carbapenemase genes (median: 0, Z=5.3, BH-adjusted p<0.001; **Table 1**). This indicates that TA systems are not preferentially located on carbapenemase plasmids compared to other AMR-encoding plasmids.

### Multi-carbapenemase genotypes have spread across many different plasmids

We identified 2447 carbapenemase plasmids from multi-carbapenemase genomes (96.3%, 1445/1501) which belonged to 307 MOB-suite secondary clusters (median number of plasmids per cluster: 2). Approximately 17.2% (422/2447) of plasmids carried more than one carbapenemase gene and were found in 133 different secondary clusters. Most plasmids carried one (43.2%, 1057/2447), two (26.3%, 643/2447) or three (16.6%, 405/2447) replicon types; the most common replicon types were IncL/M (7.5%, 183/2447), IncX3 (6.9%, 169/2447), IncN (4.8%, 118/2447), rep_cluster_1195 (ColKP3) (4.7%, 115/2447), IncX3/rep_cluster_1195 (4.5%, 110/2447), IncFIB/IncFII/rep_cluster_2183 (4.7%, 108/2447), and IncC (4.2%, 102/2447). IncF-type replicons were found on 39.8% (973/2447) of plasmids and 4.0% (99/2447) of plasmids had no replicon type. Most plasmids were predicted to be conjugative (69.4%, 1699/2447), and the remainder were predicted to be mobilizable (18.1%, 444/2447) or non-mobilizable (12.4%, 304/2447).

The top eight clusters comprised only 32.1% (785/2447) of all carbapenemase plasmids in this dataset (**Table 2, Supplementary Figure 8**). These clusters are well-known carbapenemase plasmid groups and are a mix of diverse, broad-range conjugative plasmids (e.g. AA002/AH539 [IncL/M]) in addition to plasmids restricted to single species (e.g. AA405/AI436 [IncFIB/HIB1B/rep_cluster_1254] in *K. pneumoniae*) and associated with single carbapenemase families (e.g. AC129/AL194 with OXA-232). All clusters have been observed for more than ten years and across multiple continents with one exception of cluster AA032/AH594 found solely in Asia restricted to a single *K. pneumoniae* lineage ST11. We described these clusters in more detail in the **Supplementary materials**.

**Table 2.** Features of the top eight carbapenemase plasmid clusters found in multi-carbapenemase genomes.

| MOB-suite primary/secondary cluster | Num. of plasmids | Median length (kb) | Most common replicon<sup>1</sup> | Most common predicted mobility<sup>1,2</sup> | Most common carbapenemase<sup>1</sup> | Mean num. of carbapenemase genes | Mean num. of AMR genes<sup>3</sup> | Number of genera | Num. of ST | Num. of years | Num. of countries | Num. of continents |
| --- | --- | --- | --- | --- | --- | --- | --- | --- | --- | --- | --- | --- |
| AA038/AH615 | 227 | 51.5 | IncX3,rep_cluster_1195 | conjugative | NDM-5 | 1.1 | 1.6 | 4 | 79 | 14 | 32 | 7 |
| AA002/AH539 | 125 | 64.5 | IncL/M | conjugative | OXA-48 | 1.0 | 0.5 | 4 | 52 | 14 | 21 | 6 |
| AC129/AL194 | 117 | 6.1 | rep_cluster_1195 | mobilizable | OXA-232 | 1.2 | 0 | 2 | 22 | 14 | 20 | 4 |
| AA032/AH594 | 76 | 132.4 | IncFIA/FII/R/U | non-mobilizable | KPC-2 | 1.3 | 3.9 | 2 | 5 | 10 | 2 | 1 |
| AA860/AJ272 | 65 | 154.4 | IncC | conjugative | NDM-1 | 1.2 | 7.0 | 6 | 28 | 12 | 15 | 6 |
| AA405/<br>AI436 | 60 | 347.<br>0 | IncFIB/HI1B<br>/rep_cluster<br>_1254 | conjugati<br>ve | NDM-5 | 1.1 | 12.2 | 1 | 16 | 10 | 21 | 6 |
| AA018/<br>AH561 | 58 | 118.<br>2 | IncFIB/FII/re<br>p_cluster_2<br>183 | conjugati<br>ve | KPC-2 | 1.3 | 1.7 | 3 | 16 | 14 | 12 | 6 |
| AA338/<br>AI283 | 57 | 95.1 | IncFIA/FIC | conjugati<br>ve | NDM-5 | 1.0 | 7.9 | 3 | 23 | 13 | 17 | 4 |

<sup>1</sup>Values represent the most common type and may not represent all plasmids in the cluster;
<sup>2</sup>Mobility predicted by MOB-suite based on presence of mating pair formation proteins and
relaxases; <sup>3</sup>Excludes carbapenemase genes

### Multi-carbapenemase plasmids cluster with single carbapenemase and non-carbapenemase plasmids

We clustered carbapenemase and non-carbapenemase plasmids (n=6897) using pling. Most plasmids (73.9%, 5100/6897), including all carbapenemase plasmids, belonged to a single community (based on containment index) with 744 subcommunities (DCJ-Indel distance of 4, see Methods), 248 (33.3%) of which contained carbapenemase plasmids. Singleton carbapenemase plasmids comprised almost half of these subcommunities (45.6% 113/248), highlighting the diversity of carbapenemase plasmids in this dataset. Of the remaining 135 subcommunities, 45 (33.3%, 45/135) contained both single-carbapenemase plasmids and multi-carbapenemase plasmids, and 34 of these (75.6%, 34/45) also contained additional non-carbapenemase plasmids, indicating that there is a potential network for wider mobilisation of carbapenemases in many plasmid groups.

An example of one subcommunity is shown in **Figure 3**; this subcommunity contained 17, 51, and 16 plasmids (n=84) with zero, one, and two or more carbapenemase genes respectively, and the dense network indicates very few structural changes separate this group. Most plasmids encoded a IncN-type replicon (83.1%, 66/84) and they were isolated from 6 genera (*Klebsiella* n=49, *Enterobacter* n=12, *Escherichia* n=11, *Citrobacter* n=10, *Kluyvera* n=1, and *Morganella* n=1). The most common carbapenemases were NDM-1 (19/84, 22.6%), IMP-4 (12/84, 14.3%), KPC-3 (9/84, 10.7%), KPC-2/KPC-2 (8/84, 9.5%), and KPC-2 (7/84, 8.3%).

**Figure 3.**
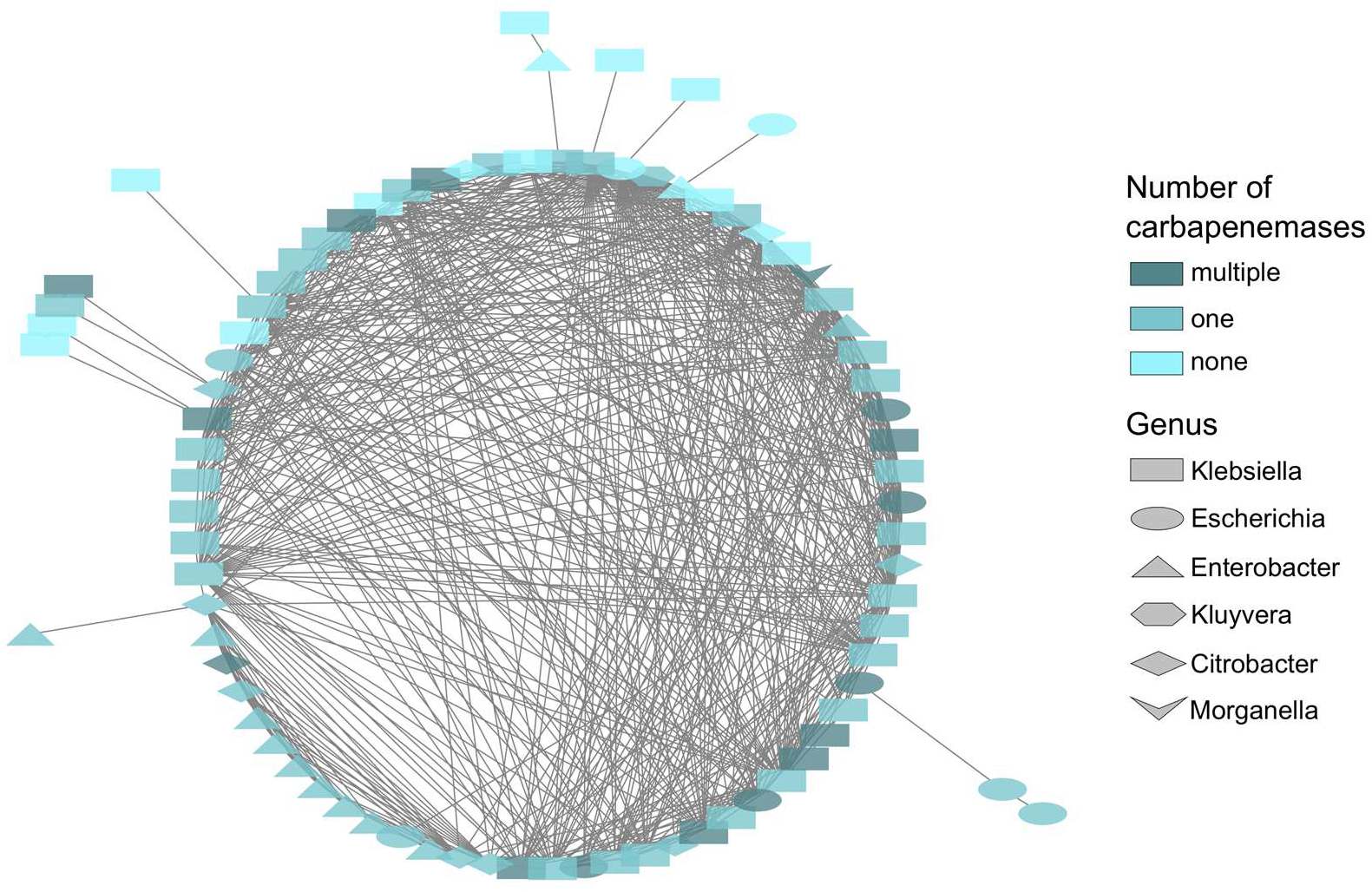
Network of one (pling) subcommunity containing plasmids encoding zero, one, or two or more carbapenemase genes. Each node represents one plasmid (n=83) coloured based on the number of encoded carbapenemase genes and shapes represent genus. Edges represent DCJ-Indel distances of 4 or smaller, indicating 4 or fewer rearrangement events separate neighbours, and edge lengths are meaningless.

### Hypervirulent multi-carbapenemase Klebsiella encode carbapenemase genes on virulence plasmids

Approximately a quarter of multi-carbapenemase *K. pneumoniae* (26.3%, 231/879) had hypervirulent genotypes (Kleborate virulence score ≥3), indicating they harboured virulence genes aerobactin, yersiniabactin, and colibactin (**Supplementary Table 8**). These genomes comprised 26 STs but were primarily ST11 (46.8%, 108/231). Most genomes were isolated from Asia (71.0%, 164/231) but at least one genome was detected from each continent. There were 53 distinct allelic carbapenemase combinations and the most common were KPC-2/KPC-2 (15.6%, 36/231), KPC-2/NDM-1 (14.7%, 34/231), NDM-1/OXA-48 (10.3%, 24/231), KPC-2/NDM-5 (8.7%, 20/231), and NDM-5/OXA-48 (8.2%, 19/231) (**Supplementary Figure 9**). These genes were found on various plasmid backbones (**Supplementary materials**). The most common aerobactin locus was *iuc1* (88.3%, 204/231), followed by *iuc5* (6.9%, 16/231), and *iuc3* (2.2%, 5/231).

Typically, hypervirulent *K. pneumoniae* encode their virulence genes on a dedicated virulence plasmid, and we examined the occurrence of carbapenemase genes on these virulence plasmids. We noted the distribution of carbapenemase types varied significantly among virulence plasmids (Pearson’s Chi-squared test, χ^2^=17.1, p<0.001); genomes with different carbapenemase families were significantly enriched in having at least one carbapenemase gene encoded on the virulence plasmid (42.1%, 61/145; OR=3.71, 95% CI [1.86, 7.81], BH-adjusted p=0.001; **Supplementary Figure 9**), whereas genomes with the same alleles rarely encoded carbapenemase genes on virulence plasmids (18.3%, 13/71; OR=0.37, 95% CI [0.16, 0.72], BH-adjusted p=0.003). Altogether, 33.5% (75/224) of virulence plasmids encoded one (92.0%, 69/75) or more (8.0%, 6/75) carbapenemase genes. These plasmids grouped into 10 secondary clusters, most of which (72.0%, 54/75) grouped in MOB-suite cluster AA405/AI436 described in **Table 2**.

### Multiple carbapenemase genes on a single plasmid backbone are located in close proximity

We examined the distance in base pairs between carbapenemase genes on plasmids that encoded more than one carbapenemase gene (n=422). Overall, the median distance between carbapenemase genes was 15.6 kb (IQR: 7.7 kb, 35.8 kb), which corresponded to a median of 18.3% (IQR: 7.6%, 33.6%) of plasmid length. Of the carbapenemase groups, plasmids with multiple NDM copies (18.6%, 78/422) had genes located the closest together, with a median of 10.5 kb apart which corresponded to a median of 7.7% of plasmid length (**Figure 4**). For multiple OXA-48-like copies (6.6%, 27/423), genes were a median of 6.1 kb apart, which corresponded to 48.4% of plasmid length; this is likely due to misassembly or duplication of small ColKP3 plasmids.

**Figure 4.**
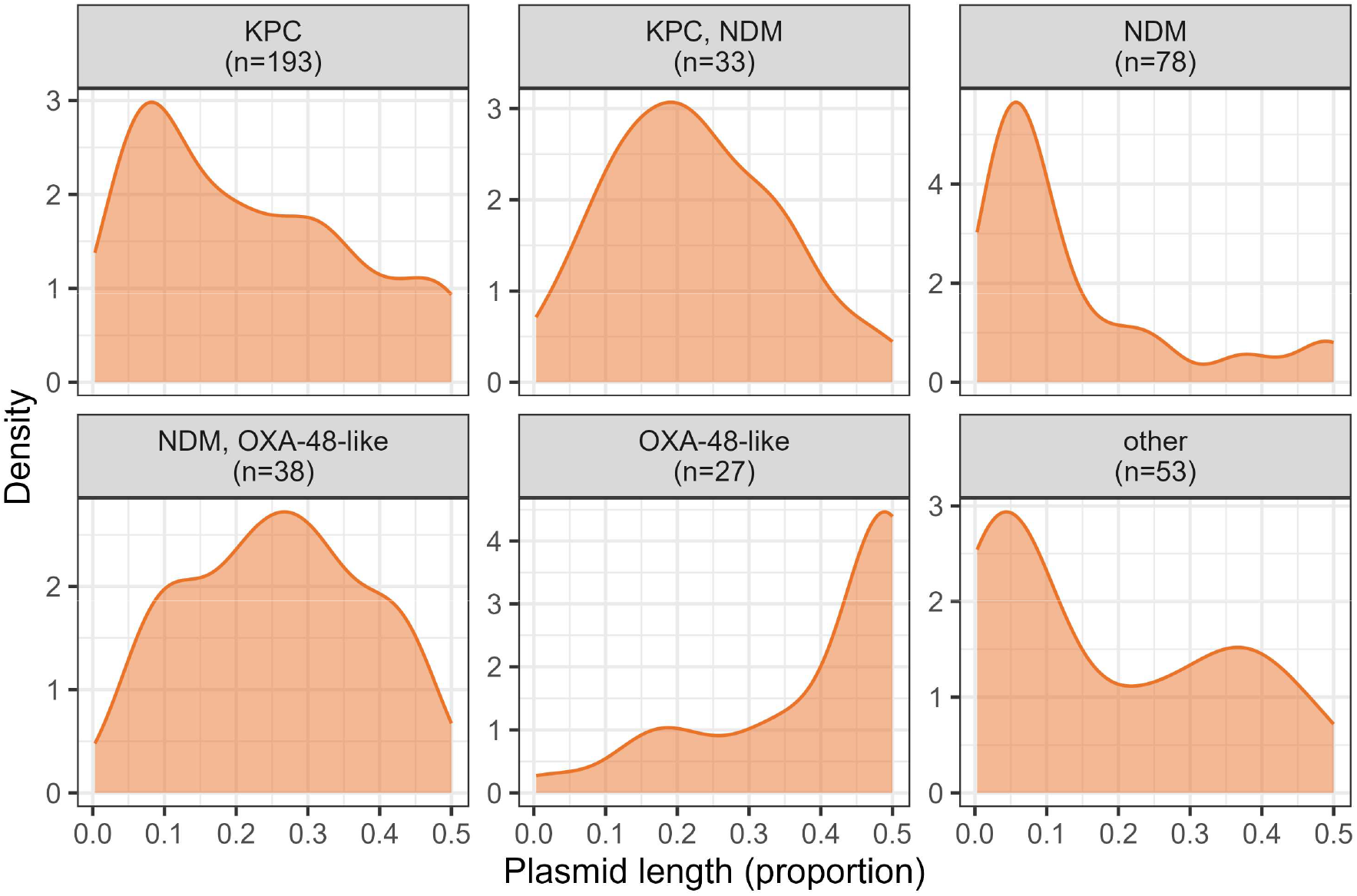
**Density of carbapenemase gene distances among proportion of plasmid length faceted by carbapenemase group**. Proportions of 0.5 indicate the second carbapenemase gene is as far away as possible i.e. on the opposite side of the circular plasmid (50 % of plasmid length).

We tested if the location of these genes on the plasmid backbones followed a uniform distribution (an equal chance of occurring anywhere on the backbone), and found that that carbapenemase groups with multiple copies of the same allele significantly deviated from uniform distributions (KPC: A^2^=9.3, p<0.001; NDM: A^2^=30.2, p<0.001; OXA-48-like: A^2^=40.8, p<0.001). However, combinations KPC/NDM and NDM/OXA-48-like did not have statistically significant deviations from uniformity (KPC/NDM: A^2^=1.6, p=0.16, NDM/OXA-48-like: A^2^=1.0, p=0.34).

### Duplication of carbapenemase genes in the same genome leads to novel gene variation

We noted that a small proportion of carbapenemase genes did not have exact allele matches to reference database sequences (4.8%, 147/3094). These were novel proteins (n=29 proteins representing n=22 novel sequences; IMP n=1, KPC n=14, NDM n=9, OXA-48-like n=5), frameshifted or pseudogenes (n=112), or had internal stop mutations (n=6) (**Supplementary Figure 10**). Only four (13.8%, 4/29) of the novel proteins appeared to have mutations in functional sites (**Supplementary Table 9**). Genes with non-exact matches were significantly enriched in genomes containing the same carbapenemase family (83.7%, 123/147) as opposed to a different family (16.3%, 24/147; 95% CI [76.7%, 89.3%], exact binomial test, p<0.001). For example, a frameshifted or novel KPC was more commonly found in a genome with other KPC as opposed to NDM or OXA-48-like. This trend was observed across all carbapenemase families and suggests that duplication of existing alleles or transposition events within a genome may facilitate novel gene variation.

### Cross-class carbapenemase gene carriage: KPC/NDM and NDM/OXA-48-like

The NDM/OXA-48-like (25.6%, 384/1501) and KPC/NDM (18.8%, 282/1501) combinations were the two most abundant combinations of different carbapenemase families and were frequently found in high-risk lineages. To evaluate the longitudinal trend of these combinations, we calculated a binary logistic regression model for each combination compared against the remaining groups (**Supplementary Figure 11, Supplementary Table 10, Supplementary Table 11**). Both models revealed a statistically significant upward trend over time (Firth’s penalised binary logistic regression; KPC/NDM: OR=1.21, 95% CI [1.15, 1.27], p<0.001; NDM/OXA-48-like: OR=1.10, 95% CI [1.06, 1.15], p<0.001), indicating that the proportion of genomes with these two combinations has increased over time relative to other combinations.

Most genomes with NDM/OXA-48-like carbapenemases carried them on two separate plasmids (78.1%, 300/384). A minority carried both NDM and OXA-48-like on the same plasmid (9.9%, 38/384) and these appeared to be increasing significantly over time (Firth’s penalised binary logistic regression, OR=1.16, 95% CI [1.02, 1.35], p=0.028) (**Supplementary Table 12**); these were grouped into 14 clusters but the most common had IncX3/rep_cluster_1195 replicons (39.5%, 15/38, MOB-suite cluster AA038/AH615) and were either NDM-5/OXA-181 (22/38, 57.9%) or NDM-1/OXA-48 (10/38, 26.3%). More information on plasmid clusters can be found in the **Supplementary materials.** A total of 38 NDM/OXA-48-like allele combinations were observed, but predominantly NDM-5 (51.6%, 198/384), NDM-1 (41.1%, 158/384), OXA-181 (38.3%, 147/384), OXA-48 (34.1%, 131/384), and OXA-232 (24.0%, 92/384). The most common combinations varied by genus, with NDM-1/OXA-48 found most frequently in *Klebsiella* (23.9%, 62/259) and *Citrobacter* (66.7%, 8/12), NDM-5/OXA-181 found most frequently in *Escherichia* (64.3%, 54/84), and NDM-5/OXA-48 found most frequently in *Enterobacter* (53.8%, 14/26).

Similar to NDM/OXA-48-like, most KPC/NDM genomes encoded the carbapenemase genes on two separate plasmids (78.7%, 222/282). A small subset carried KPC/NDM on the same plasmid (12.1%, 34/282) but there was no significant trend over time (Firth’s penalised binary logistic regression, OR=1.04, 95% CI [0.90, 1.22], p=0.56) (**Supplementary Table 13**); these grouped into 15 clusters with the most frequent being on IncL/M (18.5%, 6/34, MOB-suite secondary cluster AA002/AH528) or IncR/IncU backbones (14.8%, 6/34, MOB-suite secondary cluster AB193/AJ859). More information on plasmid clusters can be found in the **Supplementary materials.** The most commonly observed genes were KPC-2 (87.6%, 247/282), NDM-1 (73.8%, 208/282), and NDM-5 (191%, 54/282). Thirty different KPC/NDM allele combinations were observed. Across all genera, KPC-2/NDM-1 was the most common combination (*Klebsiella* (58.6%, 126/215)*, Escherichia* (41.2%, 7/17)*, Citrobacter* (64.0%, 16/25), and *Enterobacter* (41.7%, 5/12)).

## DISCUSSION

We analysed the genomic features of 1501 global genomes encoding multiple carbapenemase genes. We described two general groups of multi-carbapenemase genomes: those with different carbapenemase gene families carried on separate plasmids, and those with multiple copies of the same allele carried on the same plasmid. We found that multiple mechanisms, including plasmid co-integration, convergence with virulence, and generation of novel allele diversity through duplication, contributed to plasmid and multi-carbapenemase evolution. We demonstrated that plasmids with multiple carbapenemase genes on the same plasmid backbone had significantly higher numbers of other AMR genes compared to single and non-carbapenemase plasmids. This supports the idea that multi-carbapenemase plasmids provide selective advantages by expanding the host’s resistance profiles through acquisition of other AMR genes and virulence factors^13^, and that mobile and plastic plasmid families are more likely to acquire AMR genes and spread them^21^. The diversity of genomes and frequent representation of high-risk lineages may also be facilitating the emergence and dissemination of multi-carbapenemase genotypes.

We observed increasing proportions of NDM/OXA-48-like and KPC/NDM genomes, which agrees with other studies examining cross-class multi-carbapenemase carriage in *Klebsiella*^14,17^. It was rare to find multi-family combinations of serine-β-lactamases (i.e. KPC/OXA-48-like), indicating there are likely more advantages for a genome with a serine-β-lactamase to acquire a metallo-β-lactamases (NDM) with their broad resistance profile and frequent co-occurrence of other AMR genes^22^. Newer β-lactam/β-lactamase inhibitors (including ceftazidime/avibactam and meropenem/vaborbactam) that target serine-β-lactamases no longer function on multi-carbapenemase organisms that also encode metallo-β-lactamases (i.e. KPC/NDM)^14,23^. There is no evidence that the clinical use of newer β-lactam/β-lactamase inhibitors has driven the upward trend in KPC/NDM^23^, but this may change as more recent data becomes available.

As others have reported, carbapenemase genes are generally encoded on separate plasmids in KPC/NDM^14,23,24^ and NDM/OXA-48-like^25^ genomes, and our results show that only a minority of genomes have both carbapenemase genes co-located on the same plasmid backbone. It is still unclear what is driving the consolidation of NDM and OXA-48-like onto a single plasmid, as unlike other multi-carbapenemase plasmids, OXA-48-like plasmids typically do not encode additional AMR genes^25–27^. Consequently, it may be advantageous for them to hybridize with NDM plasmids to gain additional AMR genes. It is also unknown why the KPC-2/NDM-1 combination appeared predominantly in many different contexts, whereas NDM/OXA-48-like genomes had a larger diversity and more even spread of gene combinations. This may be related to the spread of the high-clones that carry them as well as geographic trends; KPC-2/NDM-1 and KPC-2 are generally more dominant in Asia while other combinations such as KPC-3/NDM-1 are more common in North America^14,28^ and perhaps have not yet disseminated globally.

Gene duplication is a mechanism to facilitate novel variation^29,30^, and we found that novel protein diversity arises most frequently in genomes that have copies of the same carbapenemase family. Duplication may also be a mechanism to increase copy number, which can provide an advantage by increasing resistance levels^16,31–35^. We showed that multiple copies of the same carbapenemase gene on the same plasmid backbone appear to be tandem duplications, which can be a reversible phenomenon to transiently increase AMR^36^. This duplication and evolutionary innovation are driven by associated mobile elements^30^, and tandem duplications also represent gene co-mobilisation or genetic linkage events which favour their potential for co-transfer^18,19^. However, mobile element turnover at insertion hotspots might result in the co-localisation of carbapenemase genes by random chance, as described for chromosomal defence islands^37^, and it might not necessarily be better for the host if two carbapenemase genes are co-localised. The lack of spatial relationship between different carbapenemase families on the same plasmid backbone and the higher number of replicon numbers indicate that most of these plasmids are hybridized or co-integrates, and further suggests that these genes are being co-selected, but not necessarily co-mobilised at the genetic level.

A subset of *K. pneumoniae* multi-carbapenemase genomes were hypervirulent, which others have documented^14,38–41^. Many examples of the convergence of AMR and hypervirulent plasmids through hybridisation exist^42–44^, and we demonstrated that multiple carbapenemase genes can exist in different genomic contexts in hypervirulent *K. pneumoniae*, which highlights how this convergence can arise from different genetic events. From a clinical perspective, the consolidation of multi-carbapenemase and hypervirulence onto a single plasmid is alarming and further investigation into the specific mechanisms of recombination and mobile elements involved would be useful for informing mitigation strategies.

We acknowledge that our study has several limitations from using publicly available genomes. There was a sparsity of data from certain geographical regions, which reflects a lack of resources in low- and middle-income countries where carbapenemase rates are disproportionally high^45^. Some genomes analysed are likely the result of targeted sequencing clonally-related outbreaks, which may result in over-representation of certain lineages or gene combinations. We attempted to address this by analysing a de-replicated dataset and found no difference in the observed trends. Finally, we were reliant on the quality of genome data submitted, and note that some genomes with duplicate alleles may be mis-assemblies (for example, duplication of small 6.1 kb ColKP3 plasmids encoding OXA-48-like genes was evident in this dataset and is a known problem with certain assemblers^46–48^).

Overall, we demonstrated that multi-carbapenemase Enterobacterales genomes and their gene combinations are diverse. High-risk lineages often carry multiple carbapenemase genes which has likely contributed to their global dissemination. We showed that genomes generally carry different carbapenemases families on separate plasmids, and some genomes carry multiple copies of the same carbapenemase allele which are likely to be tandem duplications when on the same backbone. Carbapenemase plasmid diversity is high in this dataset and there were many examples of multiple carbapenemase genes co-localized on the same backbone, including virulence plasmids of *K. pneumoniae*. We noted that NDM/OXA-48-like and KPC/NDM combinations are increasing in abundance and merit future monitoring. This study provides valuable insights into the genomic features of these isolates to further our understanding of their epidemiology and to inform future surveillance and treatment strategies.

## METHODS

### Identifying publicly available genomes with multiple carbapenemase genes

We obtained a dataset of isolate genome assemblies with complete chromosome and plasmid sequences containing multiple carbapenemase genes (‘multi-carbapenemase genomes’) by downloading metadata for all *Enterobacterales* (taxid: 91347) isolate assemblies in NCBI GenBank (complete, chromosome, scaffold, and contig assembly levels) using ncbi-datasets v18.29.0^49^ on 28 May 2026 (n=1278576). We did not rely solely on assemblies marked as “complete” nor those listed as generated using long-read sequencing technology as these fields are susceptible to errors during upload by individual submitters. Instead, we first applied filtering for total genome length (greater than 3 Mb and less than 7 Mb total assembly size) to remove contaminated sequences (n=1275695). Then, we kept genomes with 20 or fewer contigs (n=46531), as done previously^50^. The distribution of the number of contigs in this putatively complete subset showed that the majority (69.7%) had a long-read sequencing technology (Oxford Nanopore Technologies, PacBio) listed in their metadata, which would be consistent with generating complete and contiguous assemblies (**Supplementary Figure 12**). We ran AMRFinderPlus v4.2.7^51^ (database version: 2026-05-15.1) on this putatively complete subset of genomes using default parameters and filtered for genomes containing hits to two or more carbapenemase genes (“blaNDM|blaKPC|blaOXA-48|blaOXA-181|blaOXA-232|blaOXA-244|blaOXA-204|blaOXA-1205|blaOXA-1207|blaVIM|blaIMP”). In addition, we included an additional 23 genomes from the NEKSUS consortium^46^ which met the above criteria. The NEKSUS consortium comprises nine English NHS Trusts (groups of hospitals under the same administration) representing the largest in terms of number of emergency admissions across all seven NHS England regions. Consecutive, unselected bloodstream infection and carbapenemase-positive rectal screening isolates were collected between October 2023 and March 2024 as part of routine clinical practice. Altogether this resulted in a dataset of 1515 putatively complete isolate genomes.

In addition, AMRFinderPlus identified 17 genes that were split across contig ends (11 from PARTIAL_CONTIG_ENDX (50% < length < 90%) and 6 with BLASTX and manual inspection (length > 90%) in the Method column of output). We excluded these genes which resulted in 14/1515 (0.1%) genomes becoming single carbapenemase genomes and we eliminated these from our dataset, leaving 1501 multi-carbapenemase genomes for analysis.

### Genome dereplication

To address bias by potentially including over-sampled and genetically-related genomes from clonal outbreaks, we generated a dereplicated dataset using SKA v1.0^52^ as done previously^50^. Briefly, we analysed pairwise single nucleotide variant (SNV) distances between genomes from the same ST, and genomes were clustered into groups if they had ≤ 5 pairwise SNVs per Mb using the ‘ska distancè command and setting the SNV threshold by genus (-s 30 for *Klebsiella* (5.8 Mb mean genome size); -s 29 for *Serratia* (5.6 Mb); -s 28 for *Citrobacter* (5.5 Mb); -s 26 for *Salmonella, Escherichia,* and *Enterobacter* (5.2 Mb); -s 25 for *Providencia* (4.8 Mb); -s 22 for *Proteus* (4.2 Mb)). The genome with the highest N50 was kept as the representative from each cluster.

### High-risk lineages and hypervirulence definitions

High-risk lineages were considered to be ST101, ST11, ST14, ST147, ST15, ST16, ST1685, ST2016, ST2017, ST258, ST2502, ST273, ST307, ST392, ST512 in *K. pneumoniae*; ST10, ST1193, ST131, ST167, ST38, ST405, ST410, ST44, ST457, ST48, ST58, ST617, ST69, ST73, ST744, ST88, and ST95 in *E. coli*; ST108, ST114, ST66, and ST78 in *E. hormaechei*; ST22, ST8, and ST91 in *C. freundii*.

*K. pneumoniae* were designated as hypervirulent if genomes had a virulence score of 3 or greater from Kleborate v3.2.4 (indicating they encode the aerobactin locus, and possibly yersiniabactin and colibactin)^44^.

### Plasmid analysis

We used genomad v1.12.0^53^ to classify contigs into plasmid, chromosome, or virus, based on scores greater than 0.7, or classified into unknown if no score was >0.7 or if length was < 0.8 kb. Any contig longer than 500 kb was assumed to be chromosomal. To detect potential phage-plasmids from virus contigs encoding carbapenemase genes, we ran the tyPPing script^54^ with default settings. To assign replicon types and secondary plasmid clusters, we ran MOB-typer from MOB-suite v3.1.9^55^ with default settings (secondary cluster Mash default distance: 0.025). MOB-typer predicts plasmid mobility as conjugative (contains mating pair formation genes), mobilizable (contains a relaxase), or non-mobilizable (no detected relaxase or mating pair formation genes). To create a network of all plasmids (including both carbapenemase and non-carbapenemase), we used pling v3.0.3^56^ with default settings (--sourmash --dcj 4 --containment-distance 0.5). Pling groups plasmids into communities (containment distance > 50 % between pairs) and then further divides these into subcommunities (double cut and join indel threshold ≤ 4), while also identifying hub plasmids (not grouped into communities, but they are densely connected on the containment network and their neighbours are not interconnected). To search for toxin-antitoxin systems, we ran TAfinder2.0 with the TADB 3.0 database^57^ and included only complete systems (not orphans). AMR gene classes were obtained from the “subclass” output of AMRFinderPlus and included aminoglycoside, beta-lactam, bleomycin, colistin, fosfomycin, glycopeptide, lincosamide, macrolide, nitrofuran, phenicol, quinolone, rifamycin, streptogramin, streptothricin, tetracycline, and trimethoprim.

### Data visualization and statistical tests

Visualizations were created in R v4.6.0 using tidyverse packages^58^, patchwork v1.3.2^59^, ggpattern v1.3.1^60^, networkD3 v0.4.1^61^, treemapify v2.6.0^62^, and ComplexUpset v1.3.3^63^. The plasmid network was created in Cytoscape v3.10.4^64^.

All statistical analyses were conducted in R using an alpha of 0.05. Fisher’s exact test with Monte Carlo simulation (20000 replicates) with post-hoc analysis using adjusted standardized residuals was performed to examine genomic location. A two-sided exact binomial test was used to examine novel alleles residing in the same cell. A Chi-squared test with Monte Carlo simulation (20000 replicates) with post-hoc analysis using adjusted standardized residuals (effect size) and the Benjamini-Hochberg (BH) method to control the overall false discovery rate was used to examine the prevalence of carbapenemase combinations between genera. Carbapenemase group trends over time and continents were assessed with a multinomial logistic regression model and an ANOVA type III test. Firth’s penalised binary logistic regression which accounts for zeros in certain years was used to evaluate changing proportions of carbapenemase genomic location comparing one category against all others. Kruskal-Wallis and Dunn’s post-hoc test with BH adjustments were performed for analysed plasmid features (AMR genes, TA systems, replicons) with zero, one, or two or more carbapenemase genes. Pearson’s Chi-squared test and post-hoc Fisher’s exact test with BH adjustments were performed for the distribution of allele types among virulence plasmids and other genomic locations. The Anderson-Darling test of goodness-of-fit was used to assess if the distance between multiple carbapenemase genes on the same plasmid backbone differed from a uniform distribution between bounds of 0 and 0.5 (representing 50% of plasmid length, or the furthest possible distance on the backbone). To resolve calculation errors caused when data points exactly matched the boundary edges (0.5) due to misassemblies/duplications, the upper boundary value was adjusted to 0.500001. Pairwise gene co-occurrence was assessed using a permutation-based null model. Gene labels were randomly permuted across genomes while preserving both the number of genes per genome and the overall frequency of each gene. Pair counts were calculated using gene multiplicities within genomes; for distinct items *i* and *j*, a transaction containing *ni* and *_nj_* copies _contributed_ *_ninj_* pairs, while self-pairs contributed (^ni^). Expected pair counts were estimated as the mean across 10000 permutations. Enrichment was calculated as the ratio of observed to expected counts. Empirical one-sided p-values were calculated as the proportion of permutations with pair counts greater than or equal to the observed count, with a +1 correction. P-values were adjusted for multiple testing using the BH false discovery rate procedure. To reduce unstable estimates arising from extremely sparse combinations, only pairs observed at least 5 times were considered.

## DATA AVAILABILITY

Genome assemblies analysed in this study are available in ENA/NCBI with accession numbers listed in **Supplementary Table 1**.

## CODE AVAILABILITY

All code and metadata used in this manuscript has been deposited in a _GitHub_ repository (https://github.com/lerminin/Multi-carbapenemase-Enterobacterales), where there is also a Quarto document enabling key results and figures to be directly reproduced.

## COMPETING INTERESTS

The authors declare no competing interests.

## AUTHOR CONTRIBUTIONS

NL, DN, SL, LM, and NS conceptualised the study. NS, SL, LM, and KC acquired funding and provided supervision. NEKSUS consortium members were involved in isolate collection and processing. NL, DN, SL, LS, LM, KLH, KC, and NS contributed to writing/editing of the manuscript. All authors approved the final draft.

## FUNDING INFORMATION

This research is supported by the National Institute for Health and Care Research (NIHR) Health Protection Research Unit: Healthcare Associated Infections and Antimicrobial Resistance (NIHR207397), a partnership between the UK Health Security Agency (UKHSA) and the University of Oxford. The views expressed are those of the authors and not necessarily those of the NIHR, UKHSA or the Department of Health and Social Care.

## ACKNOWLEDGEMENTS

The authors would also like to acknowledge all participating laboratories in the NEKSUS consortium who were responsible for isolate collection, Zeynab Yusuf from UKHSA for her role in sample transportation from UKHSA to Oxford, and laboratory and bioinformatician colleagues at the Modernising Medical Microbiology Unit at the University of Oxford for support. We are grateful to Hieu Thai and to the ORACLE Corporation for access to their cloud. Computation used the Oxford Biomedical Research Computing (BMRC) facility, a joint development between the Wellcome Centre for Human Genetics and the Big Data Institute supported by Health Data Research UK and the NIHR Oxford Biomedical Research Centre. The authors would like to thank the individuals within the NEKSUS consortium group authorship are (listed alphabetically): Ian Butler, Jonathan Lambourne, Gabriella Shanks (Barts Health NHS Trust); Nicholas Brown, Poorvi Patel (Cambridge University Hospitals NHS Foundation Trust); Jasvir Nahl, Kavita Sethi (Leeds Teaching Hospitals NHS Trust); Simon Eccles (Manchester University NHS Foundation Trust); Caroline Cullerton, Julie Samuel (The Newcastle upon Tyne Hospitals NHS Foundation Trust); Nicola Childs, Samuel Stafford (North Bristol NHS Trust); Lucinda Barrett, Katie Jeffery, Sarah Oakley (Oxford University Hospitals NHS Foundation Trust); Jenny Bradbury, Christopher Jones, Jumoke Sule (UKHSA); Mark Garvey, Alan McNally (University Hospitals Birmingham NHS Foundation Trust); Martin Williams (University Hospitals Bristol and Weston NHS Foundation Trust); James Price, Paul Randell (University Hospitals Sussex NHS Foundation Trust); Gillian Rodger, Katie Hopkins, Valentina Pennetta (University of Oxford).

## Notes

https://github.com/lerminin/Multi-carbapenemase-Enterobacterales

### Competing Interest Statement

The authors have declared no competing interest.

