## Supplementary materials for "Global genomic epidemiology of multi-carbapenemase Enterobacterales"

*IMP & VIM*

A total of 110 genomes encoded IMP (110/1501, 7.3%). IMP was most commonly present in combination with NDM (44.5%, 49/110), followed by multiple IMP copies (23.6%, 26/110), and with KPC (20.9%, 23/110). There were 38 combinations detected, and the most common combination was IMP-4/NDM-1 (31.5 %, 35/110). Thirteen IMP variants were represented, with IMP-4 (57.2%, 63/110) and IMP-1 (20.0%, 22/110) comprising the majority of genomes. Most genomes encoded carbapenemase genes on two separate plasmids (64.5%, 71/110), while some had multiple carbapenemase genes on a single plasmid (27.3%, 30/110). IMP combinations were found in 7 genera and 68 STs, with *Klebsiella* (48.2%, 53/110), *Enterobacter* (27.3%, 30/110), and *Citrobacter* (14.5%, 16/110) carrying IMP combinations most frequently.

A total of 110 genomes encoded VIM (82/1501, 5.5%). VIM was most commonly found in combination with KPC (34.1 %, 28/82), followed by OXA-48-like (28.0 %, 23/82), and multiple VIM copies (19.5 %, 16/82). VIM-1 was the most common allele observed (84.1 %, 69/82), and the most common allele combinations were OXA-48/VIM-1 (23.2 %, 19/82) and KPC-2/VIM-1 (22.0 %, 18/82). As with IMP, most genomes with VIM combinations carried carbapenemase genes on two separate plasmids (64.6 %, 53/82) with a minority encoding multiple carbapenemase genes on a single plasmid (15.9 %, 13/82). VIM combinations were found in 8 genera comprising 48 STs, with *Klebsiella* (42.7 %, 35/82), *Citrobacter* (22.0 %, 18/82), and *Enterobacter* (20.7 %, 17/82) observed most frequently.

*Description of the top eight carbapenemase plasmid clusters*

The most frequent cluster was AA038/AH615 (227/2453, 9.3%) which comprised of IncX3 and IncX3/rep_cluster_1195 plasmids. These plasmids had a median size of 51.5 kb and commonly harboured OXA-181 (120/227, 52.9%), NDM-5 (72/227, 31.7%), and NDM-7 (23/227, 10.1%). This cluster contained 27 plasmids that had multiple carbapenemase genes, which were primarily NDM-5/OXA-181 (14/27, 51.9%) and NDM-5/NDM-5 (8/27, 29.6%). These plasmids were annotated as conjugative by MOB-suite and found in a diversity of species (n=10) and sequence types (n=79). The average number of AMR genes per plasmid excluding carbapenemase genes was 1.6 and other common AMR genes on these plasmids included *qnrS1* (116/227, 51.1%) and *ble* (107/227, 47.1%).

The second most common cluster was AA002/AH539 (125/2453, 5.1%) which had a median size of 64.5 kb and IncL/M replicons. These plasmids primarily harboured OXA-48 (119/125, 95.2%) across a diversity of species (n=8) and sequence types (n=52) and were annotated as conjugative. It was rare for these plasmids to harbour more than one carbapenemase (5/125, 4.0%), and those that did were primarily OXA-48-like combinations with one KPC-2/VIM-1 exception. Plasmids in this clusters carried an average of 0.5 AMR genes excluding carbapenemases, with a small subset carrying $bla_{TEM-1}$ (21/125, 16.8%) or $bla_{CTX-M-3}$ (16/125, 12.8%).

The third most prevalent cluster was AC129/AL194 (117/2453, 4.6%) which comprised of 6.1 kb plasmids with rep_cluster_1195 or ColKP3 replicons. OXA-232 (109/117, 93.2%) or other OXA-48-like alleles were found in this cluster, and these plasmids were predicted to be mobilizable due to encoding a MOBP relaxase. These plasmids were mainly found in *K. pneumoniae* and several *E. coli* genomes across various STs. Plasmid size scaled with the number of carbapenemase genes; 19

plasmids doubled in size (12 kb) had two OXA-48-like copies, and two plasmids tripled in size (18 kb) had three OXA-48-like copies. These are likely assembly duplication errors as opposed to true biological duplication which is a known issue for small plasmids assembled with certain long-read assemblers[4–6]. No other AMR genes were found on these plasmids.

Cluster AA032/AH594 (76/2453, 3.1%) had a median size of 132.4 kb and most plasmids encoded KPC-2 (68/76, 89.5%). These plasmids were had IncFIA/IncFII/IncR/IncU replicons (42/76, 55.3%) or variations, and the majority were predicted to be mobilizable (48/76, 63.2%), whereas some were predicted to be non-mobilizable (17/76, 22.4%) and some conjugative (11/76, 14.5%). Most plasmids were found in *K. pneumoniae* ST11 (69/76, 90.8%), with a few found in other STs (6/76, 7.9%) and one found in *Morganella morganii* ST8 that had an additional IncN replicon. All but one plasmid was isolated from China from 2017 to 2025. This cluster contained 23 plasmids that encoded multiple carbapenemases, where multiple KPC-2 copies was most common (12/23, 52.2%), followed by KPC-2/NDM-1 (4/23, 17.4%), KPC-2/NDM-5 (4/23, 17.4%), one KPC-2 with two novel NDM matches, one KPC-14/KPC-14, and one IMP-4/KPC-2. These plasmids carried a mean number of 3.9 AMR genes excluding carbapenemase genes, with *bla*$_{\text{TEM-1}}$ (55/76, 72.4%), *rmtB1* (55/76, 72.4%), *bla*$_{\text{CTX-M-65}}$ (54/76, 71.1%), and *bla*$_{\text{SHV-12}}$ (53/76, 69.7%) commonly found on these backbones.

Cluster AA806/AJ272 (65/2453, 2.6%) encoded IncC (48/65, 73.8%) or IncC/rep_cluster_1527 (11/65, 16.9%) replicons or other variations and had a median size of 154.4 kb. This cluster had all 5 carbapenemase families represented, with NDM-1 (46/66, 69.7%) being the most common, followed by OXA-181 (8/65, 12.3%), other NDM alleles (10/65, 15.4%), KPC-2 (2/65, 3.1%), and one KPC-3, IMP-26, OXA-48, and VIM-4, respectively. This cluster was found in the widest breadth of genera (n=6) representing 28 STs. Ten plasmids encoded multiple carbapenemases, with multiple NDM copies being the most prevalent (6/10, 60.0%) and one IMP-26/VIM-4, one KPC-2/NDM-1, one NDM-1/OXA-181, and one novel NDM/OXA-48. Plasmids in this cluster carried an average of 7 AMR genes excluding carbapenemase genes and other genes such as *sul1* (62/65, 95.4%), *ble* (55/65, 84.6%), *aac(6')-Ib3* (33/65, 50.8%), *rmtC* (32/65, 49.2%), and *bla*$_{\text{CMY-6}}$ (28/65, 43.1%).

Cluster AA405/AI436 (60/2453, 2.4%) plasmids had the largest median size of 347.0 kb and encoded replicons IncFIB/IncHI1B/rep_cluster_1254 (25/60, 41.7%) or IncFIB/IncHI1B (18/60, 30.0%) or other variations. These plasmids encoded NDM-5 (25/60, 41.7%), NDM-1 (23/60, 38.2%), OXA-48 (11/60, 18.3%), OXA-232 (3/60, 5.0%) or KPC-2 (1/60, 1.7%). This cluster was predicted to be conjugative and only found in *K. pneumoniae* across multiple STs (n=16), with ST147 (14/60, 23.3%) and ST383 (10/60, 16.7%) being the most prevalent. Four plasmids contained multiple carbapenemases: three NDM-1/OXA-48 and one with three copies of NDM-5. These plasmids contained the highest average number of AMR genes excluding carbapenemase genes per plasmids at 12.2, with *sul1* (55/60, 91.7%), *ble* (48/60, 80.0%), *qnrS1* (47/60, 78.3%), *aph(3')-VI* (45/60, 75.0%), *mph(A)* (44/60, 73.3%), *mrx(A)* (44/60, 73.3%), *dfrA5* (43/60, 71.7%), *bla*$_{\text{CTX-M-15}}$ (38/60, 63.3%), *catA1* (35/60, 58.3%), and *bla*$_{\text{TEM-1}}$ (33/60, 55.0%) being the most common genes. Notably, 48/60 (80.0%) plasmids in this cluster also contained virulence genes for aerobactin, yersiniabactin, and colibactin.

Cluster AA018/AH561 (58/2453, 2.4%) had IncFIB/IncFII/rep_cluster_2183 replicons, predicted to be conjugative, and were a median 118.2 kb in size. This cluster exclusively encoded KPC carbapenemases, with KPC-2 (25/58, 43.1%) and KPC-3 (26/58, 44.8%) being the most frequent. Nineteen plasmids had duplicate KPC copies, most of which were KPC-3/KPC-3 (11/19, 57.9%). These plasmids were mainly found in *K. pneumoniae* in multiple lineages, notably in ST512 (20/58, 34.5%) and ST258 (9/58, 15.5%), with three plasmids were found outside *K. pneumoniae* in *E. coli*, *Klebsiella ornithinolytica*, and *C. freundii*. Plasmids in this cluster had an average of 1.7 AMR genes per plasmid, with *bla*$_{\text{OXA-9}}$ (42/58, 82.8%) and *bla*$_{\text{TEM-1}}$ (39/58, 67.2%) being the most common genes in addition to carbapenemase genes.

Plasmids in cluster AA338/AI283 (57/2453, 2.3%) were predicted to be conjugative, were 95.1 kb in size and had IncFIA/IncFIC (39/57, 68.4%) or IncFIA (10/57, 17.5%) or other hybrid replicons. NDM-5 was the most common carbapenemase in this cluster (52/57, 91.2%), in additional to NDM-1 (2/57, 3.5%) and novel NDM (3/57, 5.3%). Plasmids in this cluster did not carry multiple carbapenemase genes. This cluster was found in many *K. pneumoniae*, several *E. coli*, and one *E. hormaechei*, altogether comprising 23 STs. The average number of AMR genes per plasmid was 7.9, with *ble* (54/57, 94.7%), *sul1* (54/57, 94.7%), *dfrA12* (52/57, 91.2%), *aadA2* (50/57, 87.7%), *bla*$_{\text{TEM-1}}$ (41/57, 71.9%), *erm(B)* (40/57, 70.2%), *mph(A)* (40/57, 70.2%), *mrx(A)* (40/57, 70.2%), and *rmtB1* (40/57, 70.2%) found commonly in this cluster.

*Carbapenemase plasmid clusters found in genomes with NDM/OXA*-48-like

Genomes encoding NDM-5 as part of the NDM/OXA-48-like combination had 33 distinct NDM-5 plasmid clusters, with the most common backbones being IncFIA/IncFIC (47/190, 24.7%, cluster AA338/AI283, see above), IncX3/rep_cluster_1185 (34/190, 17.9%, cluster AA038/AH615, see above), IncFIB/IncHI1B/rep_cluster_1254 (20/190, 10.5%, cluster AA405/AI436 with virulence genes, see above), and IncFIA/IncFIB/IncFII/IncFII/IncQ1 (16/190, 8.4%, cluster AH831). NDM-1 was found on 40 distinct clusters in the NDM/OXA-48-like isolates, with IncFIB/IncHI1B/IncR/rep_cluster_1254 (28/153, 18.3%, cluster AA405/AI439 with virulence genes), IncC (19/153, 12.4%, cluster AA806/AJ272, see above), and IncFIB/IncHI1B/rep_cluster_1254 backbones (17/153, 11.1%, cluster AA405/AI436 with virulence genes, see above). OXA-48-like alleles were found in fewer background than NDM in NDM/OXA-48-like genomes; OXA-181 was found in 13 distinct clusters with the most frequent being IncX3/rep_cluster_1195 backbones (106/136, 77.9%, cluster AA038/AH615, see above), OXA-48 was found in 16 clusters with IncL/M backbones (78/127, 61.4%, cluster AA002/AH539, see above; 23/127, 18.1%, cluster AA002/AH529) being the most common, and OXA-232 was found almost exclusively on ColKP3/rep_cluster_1195 backbones (92/93, 98.9%, cluster AC129/AL194, see above).

*Carbapenemase plasmid clusters found in genomes with KPC/NDM*

Isolates encoding KPC-2 as part of the KPC/NDM combinations harboured the gene on 60 different plasmid clusters, mostly commonly on IncFIA/IncFII/IncR/IncU (46/247, 18.6%, cluster AA032/AH594 see above) and IncFIA/IncFII/IncR backbones (30/247, 20.4%, cluster AA032/AH591). Similarly, genomes encoding NDM-1 as part of the KPC/NDM combination had 47 different NDM-1 plasmid clusters, primarily IncFIB/IncFII/rep_cluster_2272 (31/211, 14.7%, cluster AA046/AH632), IncX3 (27/211, 12.8%, cluster AA038/AH618), and IncU/rep_cluster_1254 backbones (19/211, 9.0%, cluster AC212/AL310).

*Plasmid clusters found in hypervirulent* Klebsiella *genomes*

There were 53 distinct allelic carbapenemase combinations in hypervirulent *K. pneumoniae* and the most common were KPC-2/KPC-2 (15.6%, 36/231), KPC-2/NDM-1 (14.7%, 34/231), NDM-1/OXA-48 (10.3%, 24/231), KPC-2/NDM-5 (8.7%, 20/231), and NDM-5/OXA-48 (8.2%, 19/231).

Plasmids with KPC-2 was found on 18 distinct clusters, with IncFIA/IncFII/IncR/IncU backbones (55/128, 43.0%, cluster AA032/AH594, see above), IncFIA/IncFII backbones (19/128, 14.8%, cluster AC037/AL057), and IncFIA/IncFII/IncR backbones (16/128, 12.5%, cluster AA032/AH591) being the most common. NDM-1 and NDM-5 were found on 32 and 21 distinct plasmid clusters respectively, with IncFIB/IncHI1B/rep_cluster_1254 backbones found more frequently in both groups (19/92, 120.6%, and 23/59, 39.0%, respectively, cluster AA405/AI436, see above).

Eighty-one plasmids encoded more than one carbapenemase and they grouped in 19 clusters. The most common clusters had IncFIA/IncFII/IncR/IncU replicons (17/81, 21.0%, cluster AA032/AH594) with KPC-2/KPC-2 (10/17, 58.8%), IncFIA/IncFII replicons (17/81, 21.0%, cluster AC037/AL057) with KPC-2/KPC-2 (9/17, 52.9%), and rep_cluster_1195 replicons (13/81, 16.0%, cluster AC129/AL194) with OXA-232/OXA-232 (6/13, 46.2%).

*Novel protein sequences*

We investigated the sequences of the novel proteins that did not have exact allele matches (n=29; IMP n=1, KPC n=14, NDM n=9, OXA-48-like n=5) (**Supplementary Figure 10**). Only four (13.8%) of the novel proteins appeared to have mutations in functional sites (**Supplementary Table 9**). Three of these were OXA-48-like proteins with OXA-232 as the closest variant; two had T213A and R214S mutations in the active site and one has R214S only[1]. The final one was an NDM variant with NDM-1 as the closest allele match but had an A233P mutation in an active site[2] along with other adjacent substitutions (A231S, A235R, and A239T). None of the novel KPC proteins appeared to have mutations in the active sites[3].

We searched for synonymous mutations and found 13 carbapenemase genes (0.4%, 13/3094; IMP-1 n=7, KPC-2 n=4, KPC-3 n=1, VIM-1 n=1) with variation in their nucleotide sequences from 9 genomes. Only one of these genomes contained two distinct nucleotide sequences that coded for the same protein (KPC-2).
