## Supplementary Figure for "Global genomic epidemiology of multi-carbapenemase Enterobacterales"

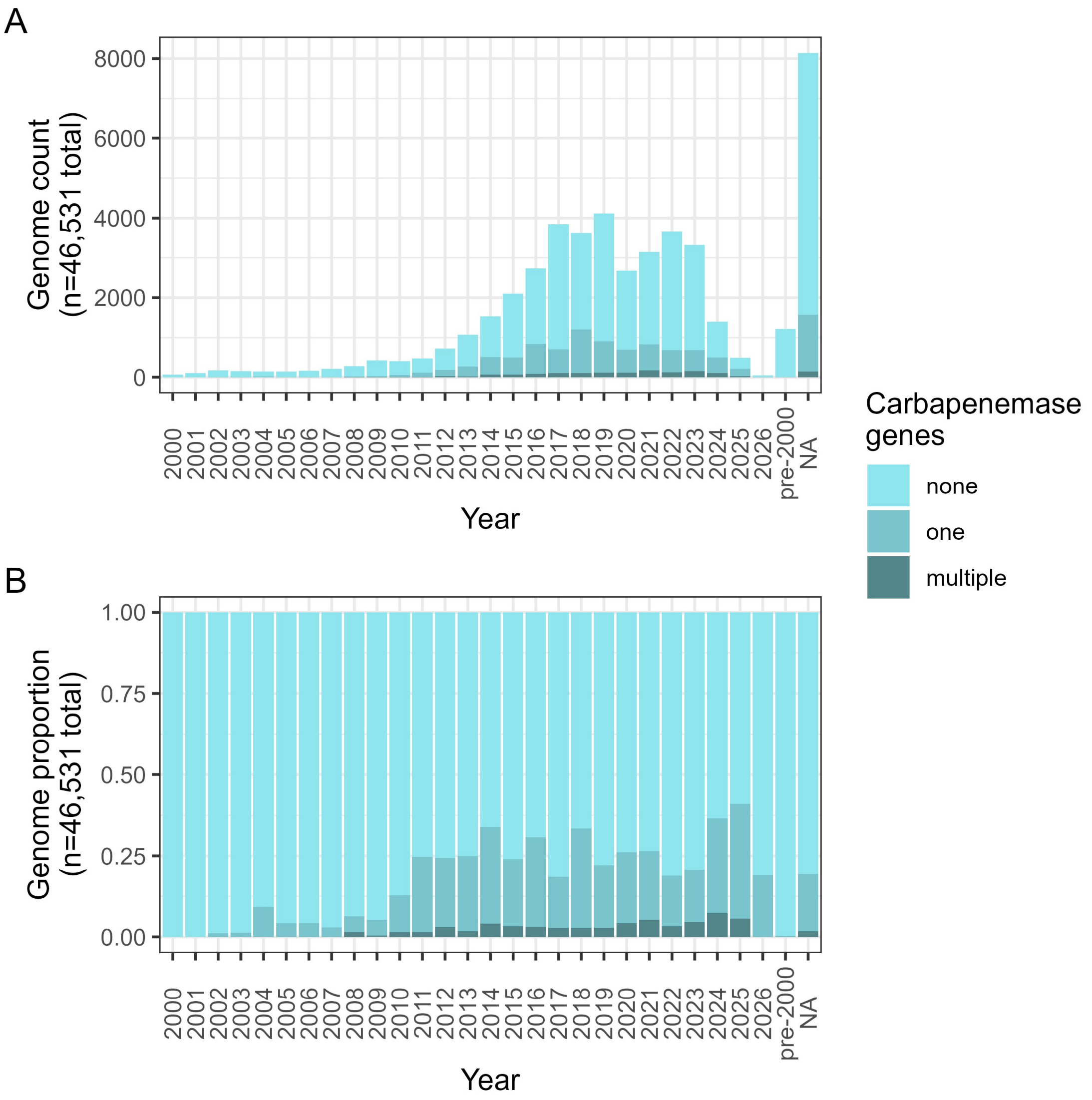


**Supplementary Figure 1. Proportion of all Enterobacterales genomes in NCBI (≤ 20 contigs) that encoded zero, one, or multiple carbapenemase genes. (A)** Total count and **(B)** proportion of 46531 genomes. “Multi” indicates two or more carbapenemase genes.

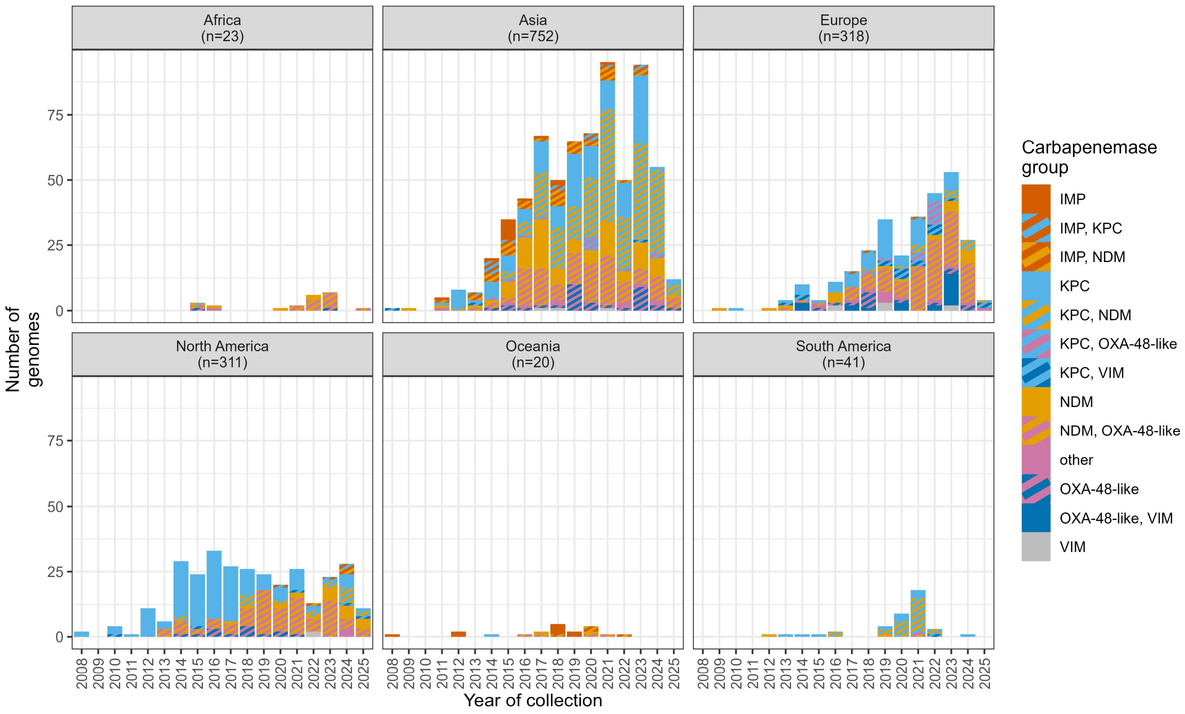


**Supplementary Figure 2.** Count of carbapenemase combinations by continent over time. “Other” includes NDM/VIM (n=8), IMP/OXA-48-like (n=4), IMP/VIM (n=3), IMP/KPC/NDM (n=5), KPC/NDM/OXA-48-like (n=3), KPC/NDM/VIM (n=2), KPC/OXA-48-like /VIM (n=1), NDM/OXA-48-like /VIM (n=1).

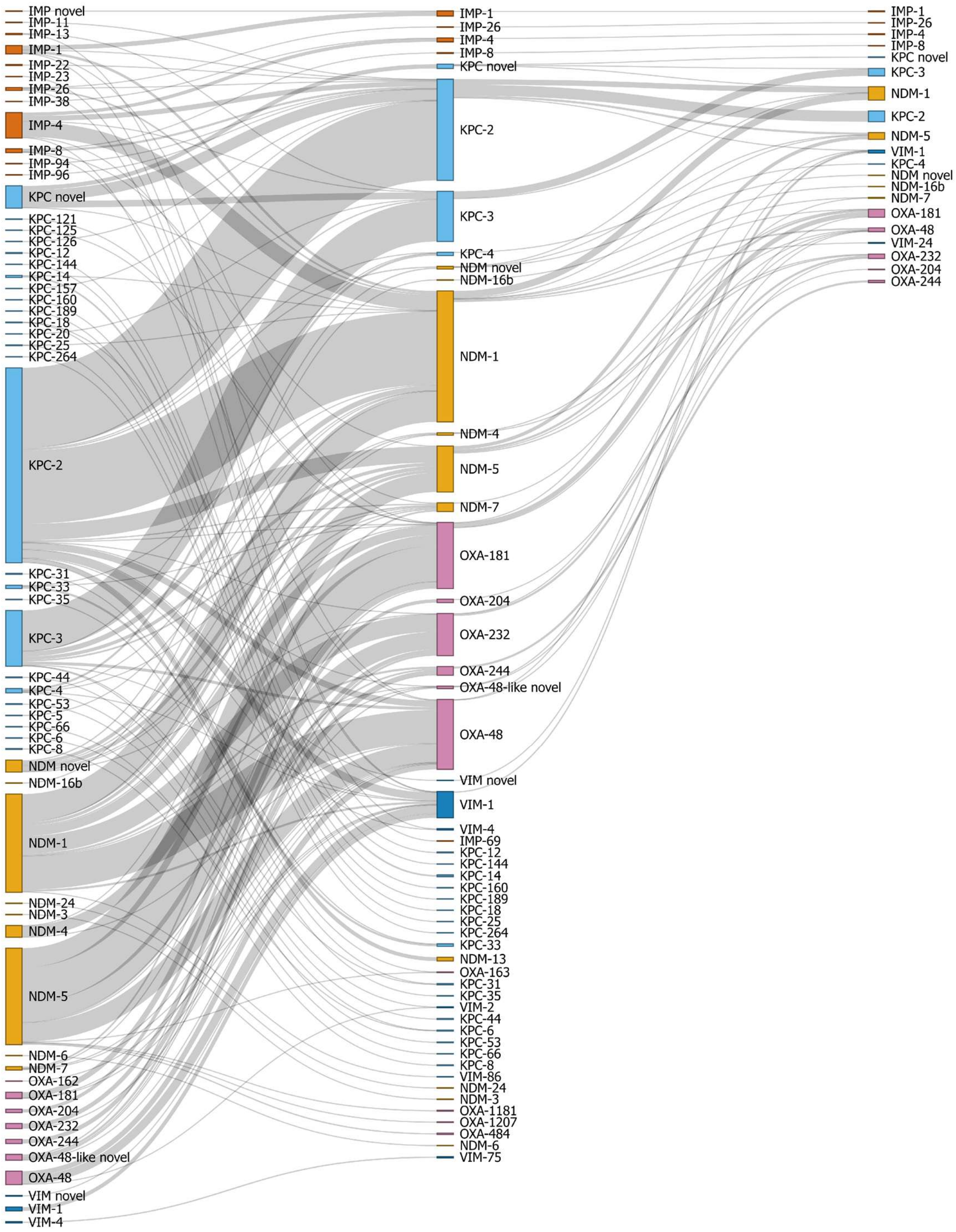


**Supplementary Figure 3**. **Linkage between carbapenemase allele combinations**. Isolates with four or more carbapenemase genes (3.5%, 53/1501) were excluded from this plot. Nodes are coloured by carbapenemase family and are ordered alphabetically. Genes labelled "novel" indicate novel protein sequences that do not match existing alleles and are not necessarily all the same sequence.

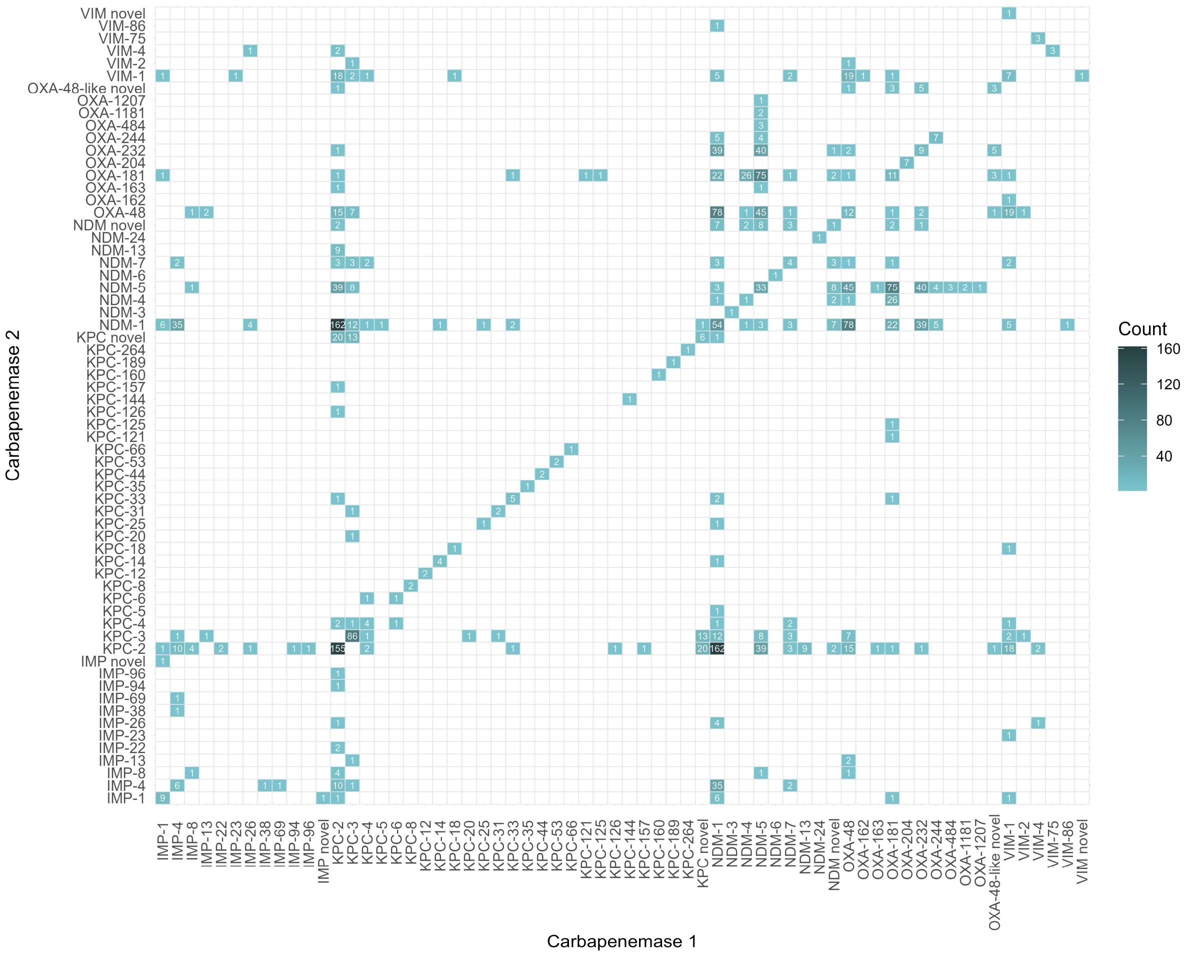


**Supplementary Figure 4**. **Heatmap of carbapenemase allele combinations (n=140/241) for genomes with two carbapenemases (86.0 %, 1290/1501)**. Darker colours indicate more genomes, and white indicates no genomes. Genomes with three or more carbapenemase genes (14.1 %, 211/1501) were not plotted (101 combinations). Proteins labelled "novel" indicate novel protein sequences that do not match existing alleles and are not necessarily all the same sequence. Note that three alleles (OXA-163 n=2, KPC-12 n=2, KPC-189 n=1) are annotated as extended-spectrum β-lactamases in the Beta-Lactamase DataBase (http://bldb.eu/BLDB.php) and display weak carbapenemase activity (https://doi.org/10.1128/aac.00022-11, https://doi.org/10.1007/s10096-021-04198-2, https://doi.org/10.1128/msystems.00722-24).

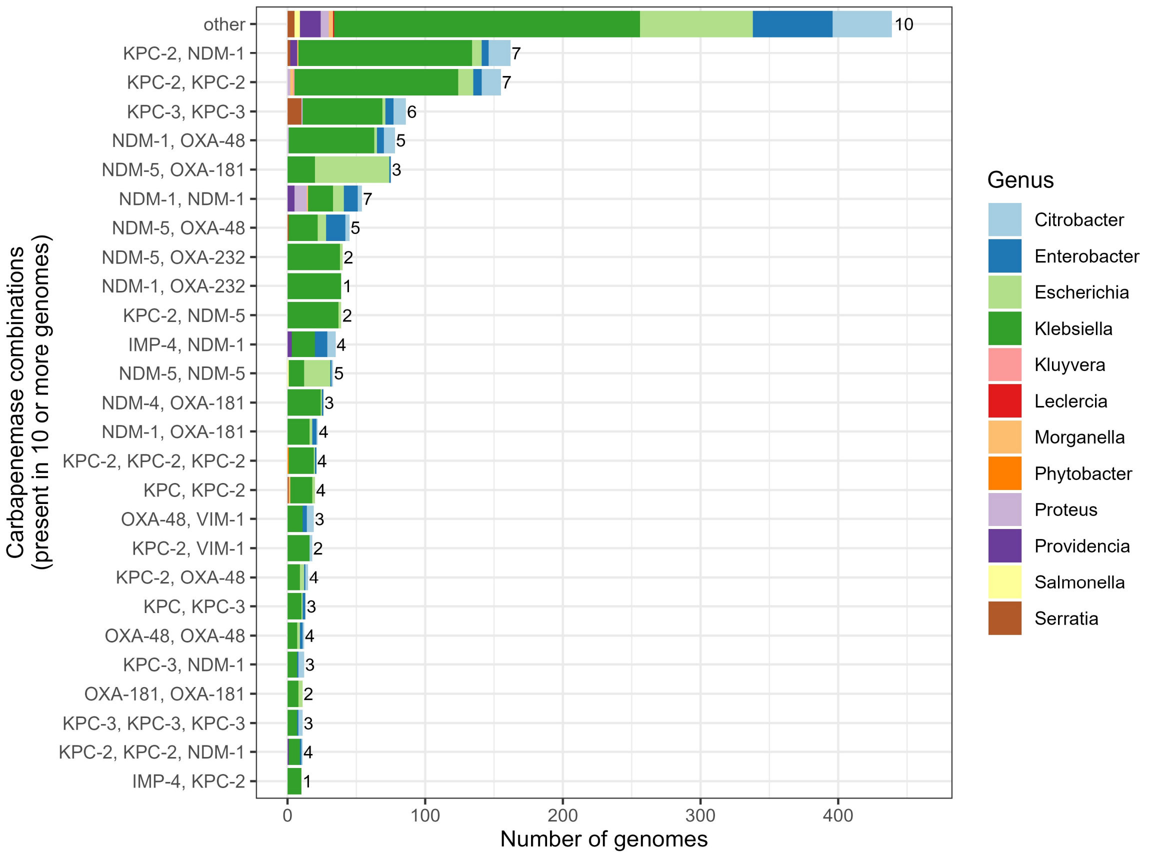


**Supplementary Figure 5. Most common carbapenemase allelic combinations found across different genera**. Combinations present in fewer than 10 genomes were grouped into “other” (n=215 combinations from 439 genomes). Numbers to the right of columns indicate the number of distinct genera where this combination was observed.

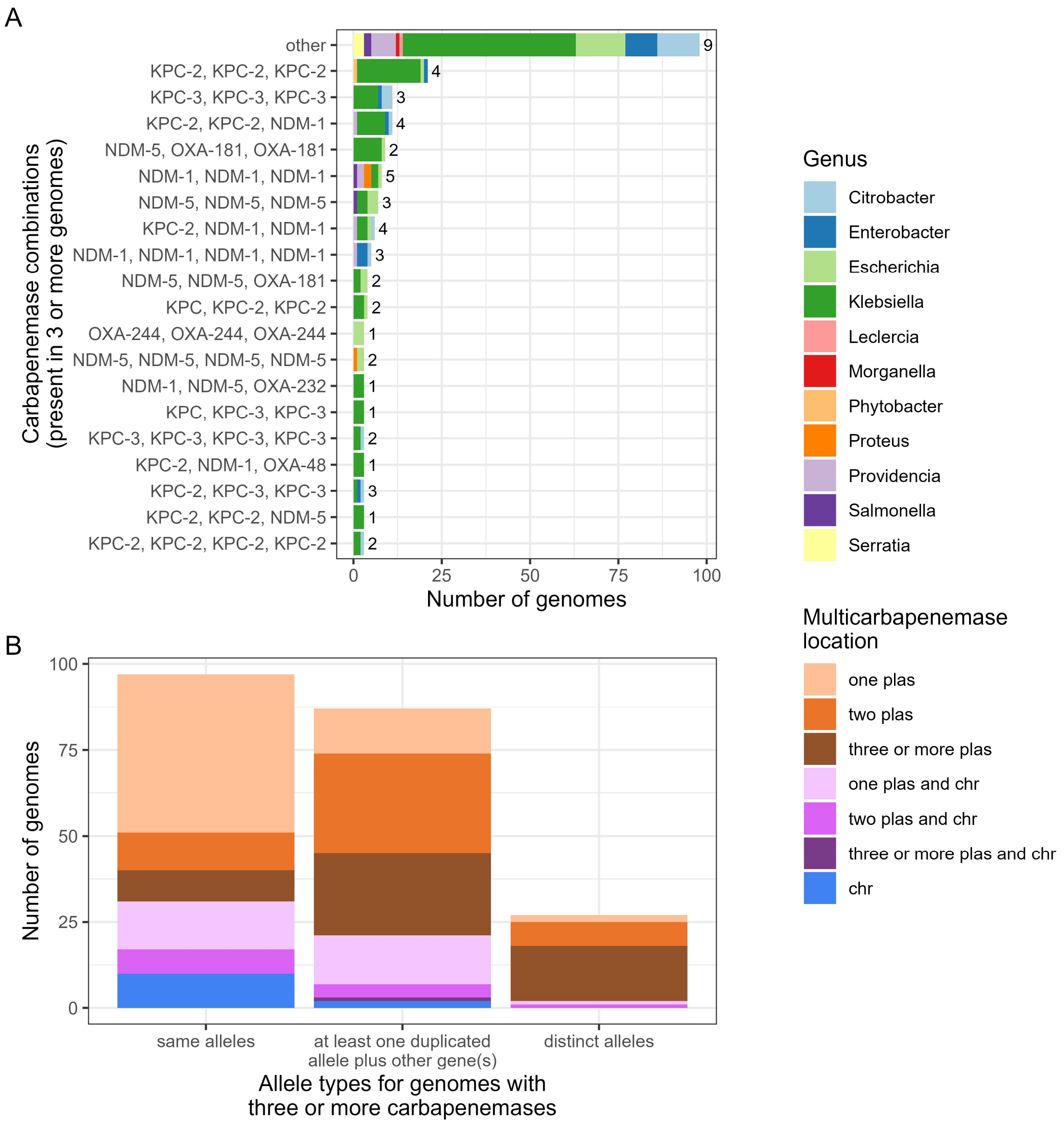


**Supplementary Figure 6.** (**A**) **Most common allele combinations in genomes with three or more carbapenemases coloured by genus (n=211)**. "Other" includes 82 additional allele combinations only found in two or fewer genomes (n=98). (**B**) **Genomic location of carbapenemase genes in genomes with three or more carbapenemases**. "Distinct alleles" includes different gene alleles (KPC-2 vs KPC-3) as well as different gene families (KPC vs NDM).

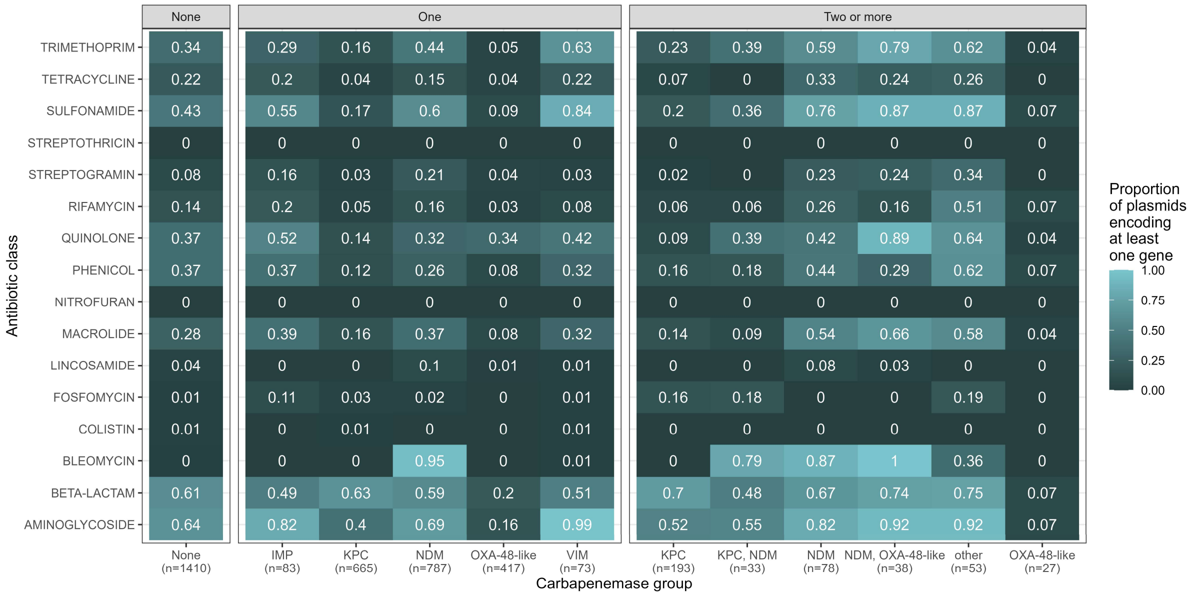


**Supplementary Figure 7. Proportion of AMR-encoding plasmids that encoded at least one resistance gene to different antibiotic gene classes.** Facets indicate plasmids with zero ("None", n=1410), one ("One", n=2025) or multiple ("Two or more", n=422) carbapenemase genes. "Other (n=53)" indicates carbapenemase groups present on fewer than 15 plasmids (IMP n=13, IMP/KPC n=6, IMP/NDM n=14, IMP/VIM n=2, KPC/NDM/VIM n=2, KPC/OXA, n=4, KPC/VIM n=3, NDM/VIM n=3, VIM n=6). Carbapenemase genes are not included in the BETA-LACTAM category.

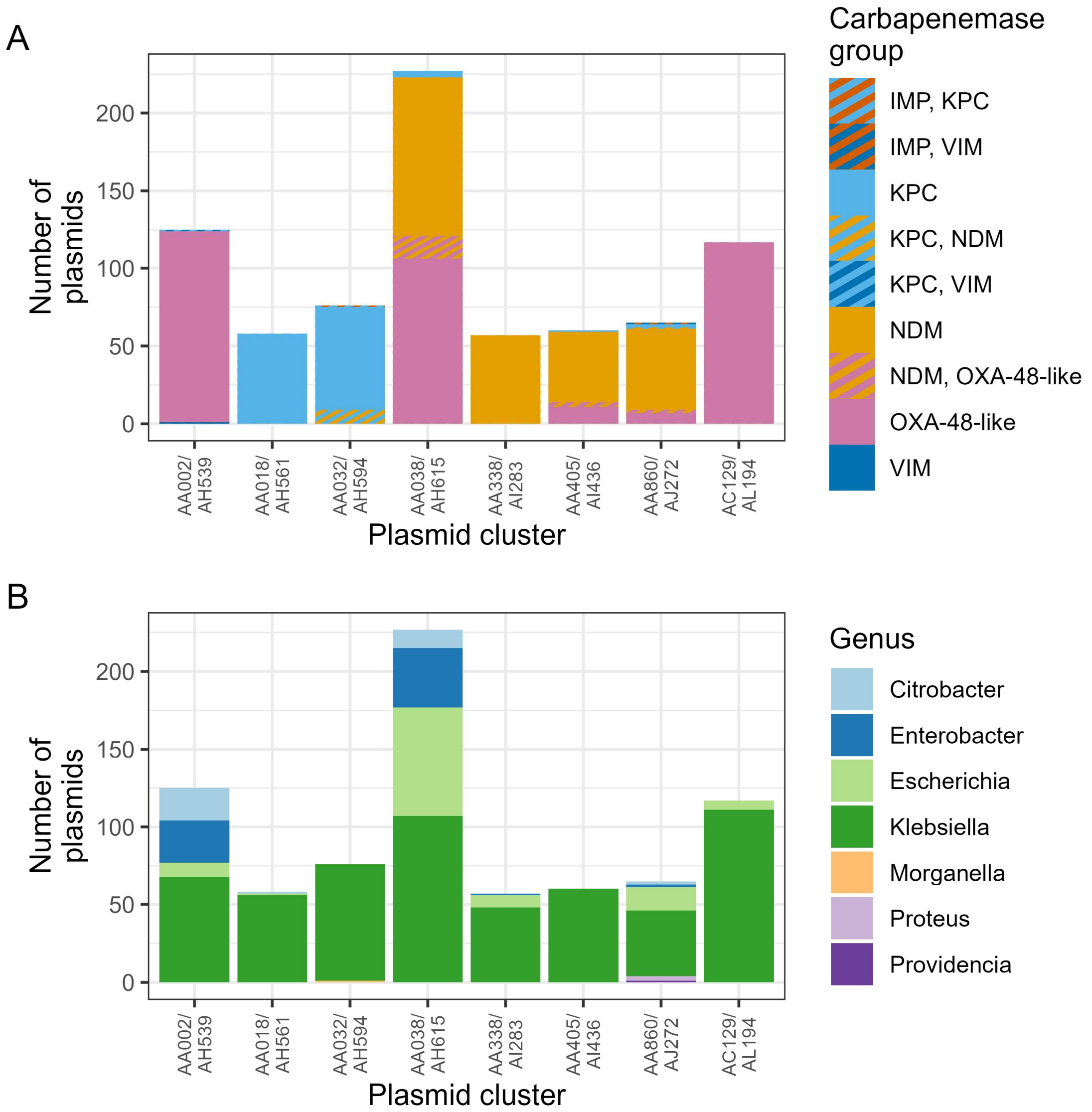


**Supplementary Figure 8. (A) Carbapenemase groups and (B) genera represented in the top 8 carbapenemase plasmid secondary clusters.** Cluster codes are secondary clusters from the default MOB-suite database.

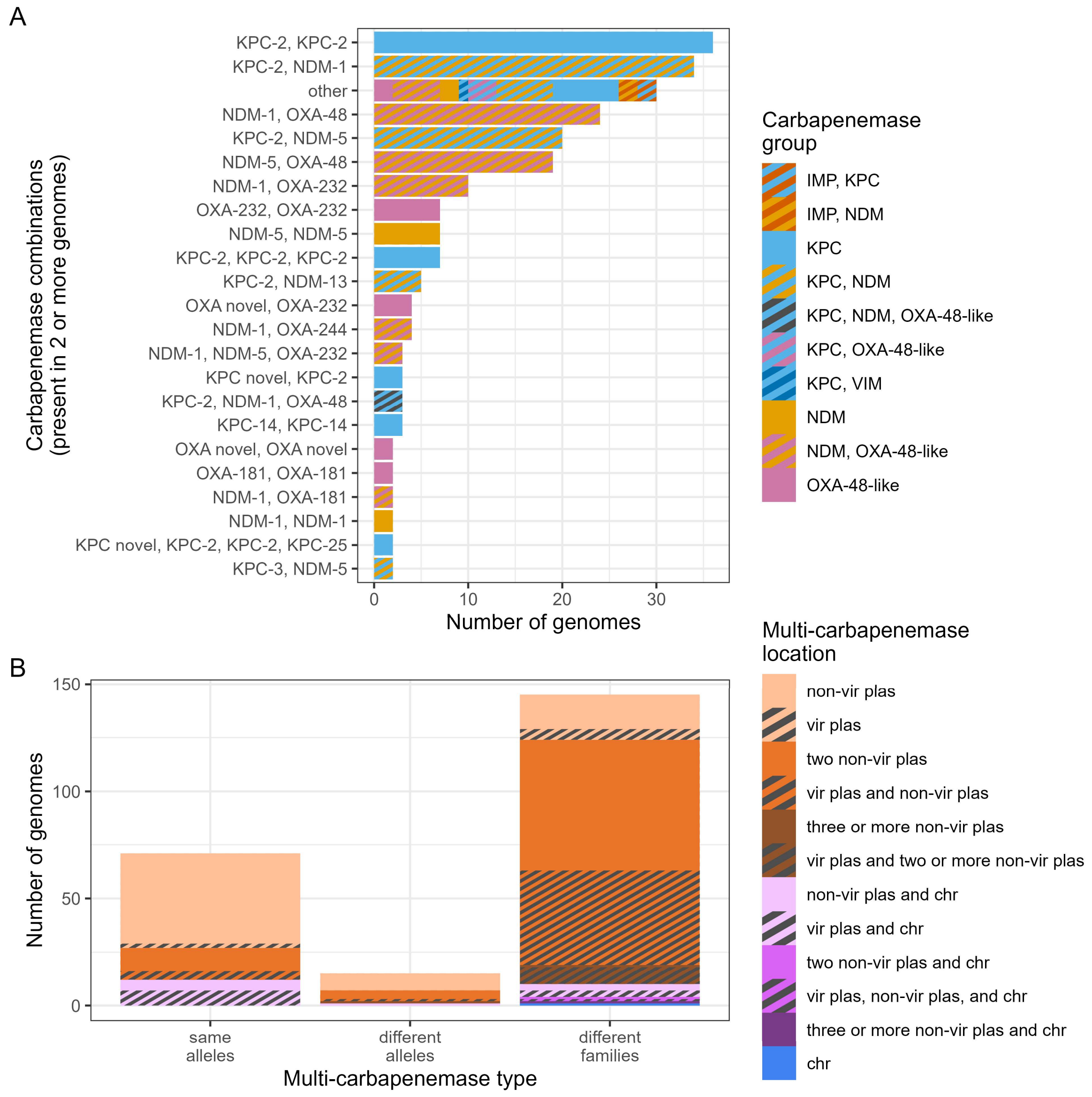


**Supplementary Figure 9.** (**A**) **Count of common carbapenemase alleles among hypervirulent *K. pneumoniae***. 30 singleton multi-carbapenemase combinations were grouped into "other". (**B**) **Genomic location of carbapenemase genes in hypervirulent *K. pneumoniae***. Black hatches represent genomes with one or more carbapenemase genes located on a virulence plasmid which carries the aerobactin locus. "plas" = plasmid, "chr" = chromosome, "vir" = virulence.

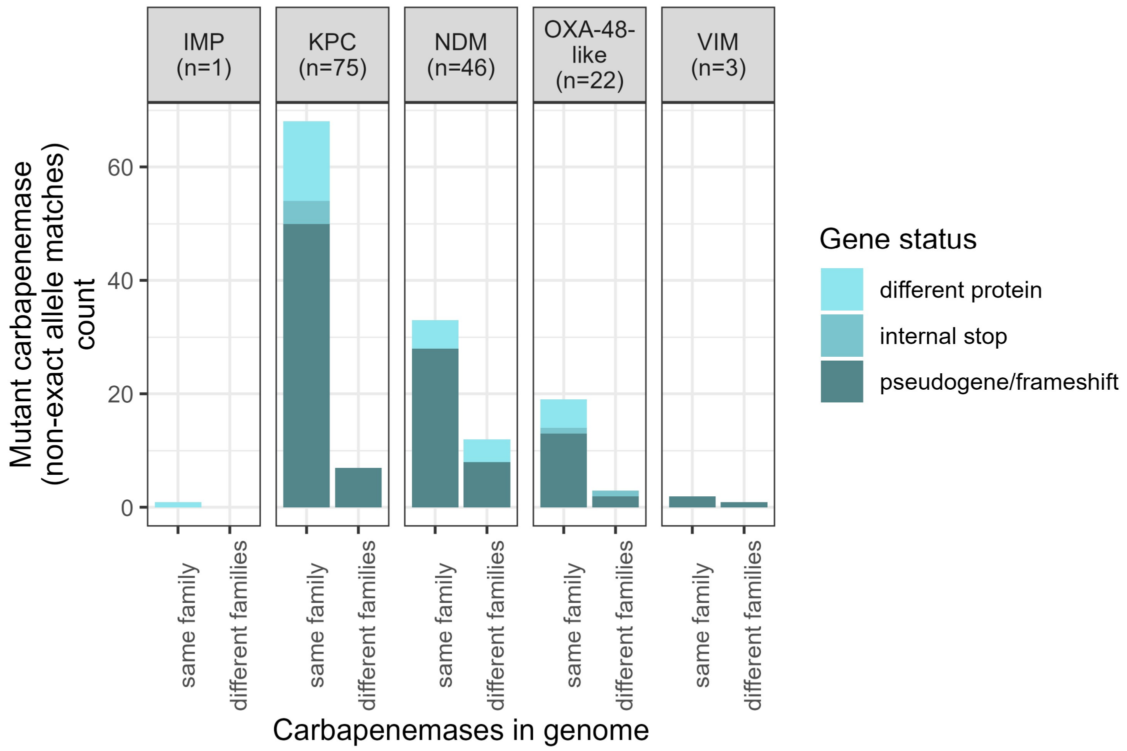


**Supplementary Figure 10**. Carbapenemases with non-exact allele matches (n=147) are more frequently found in genomes harbouring the same family of carbapenemase as opposed to different families. "Same family" indicates the non-exact allele was found in the same genome with another gene of the same carbapenemase family (i.e. IMP/KPC/NDM/OXA-48-like/VIM), whereas "different families" indicate a different carbapenemase gene family was found in the same genome. Gene status was obtained from the "Method" column in the AMRFinderPlus output, where "different protein" = BLASTX (BLAST alignment is >90% of length and >90% identity to a protein in the AMRFinderPlus database), "internal stop" = INTERNAL_STOP (translated blast reveals a stop codon that occurred before the end of the protein), and "pseudogene/frameshift" = PARTIALX (BLAST alignment is >50% of length, but <90% of length and >90% identity to the reference, and does not end at a contig boundary).

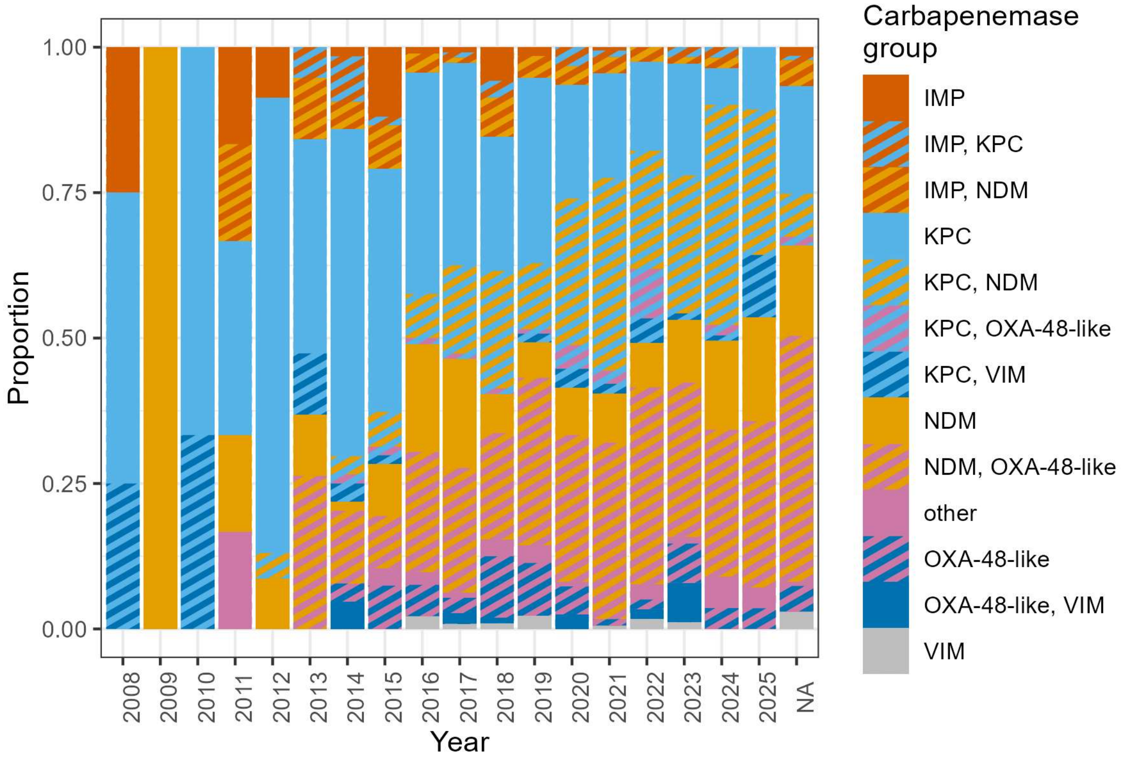


**Supplementary Figure 11. Proportion of genomes with different multi-carbapenemase groups over time.** "NA" indicates year of collection was not included in the assembly metadata. "Other" includes NDM/VIM (n=8), IMP/OXA (n=4), IMP/VIM (n=3), IMP/KPC/NDM (n=5), KPC/NDM/OXA (n=3), KPC/NDM/VIM (n=2), KPC/OXA/VIM (n=1), NDM/OXA/VIM (n=1).

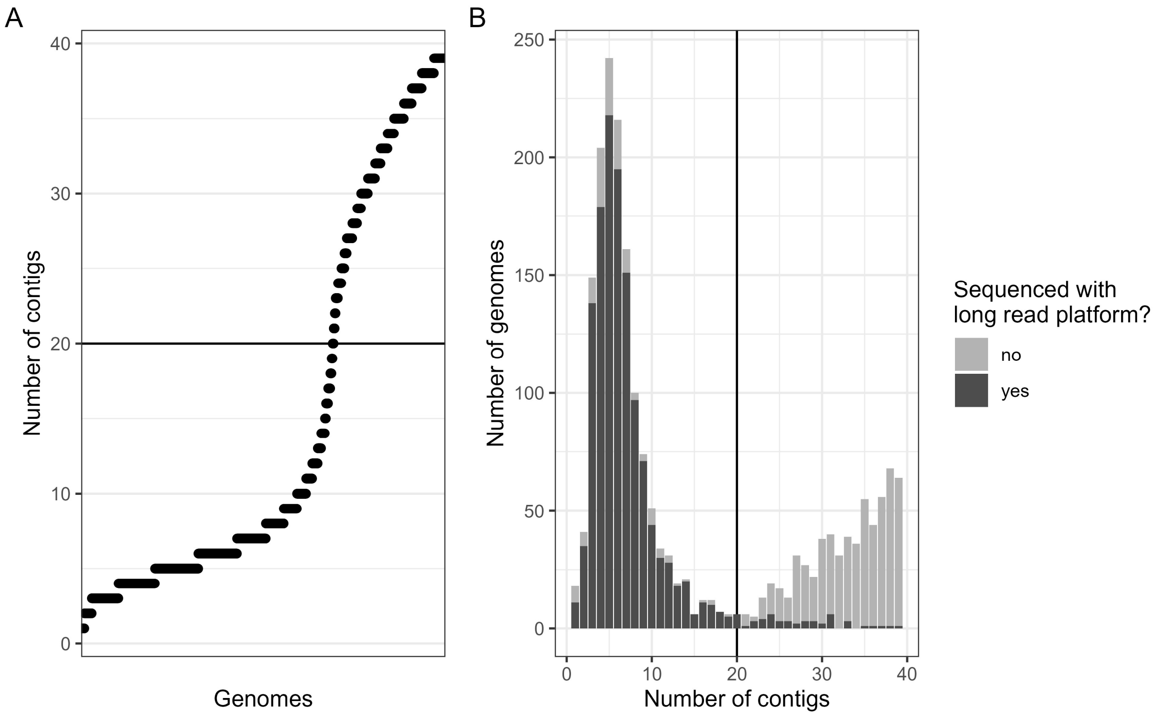


**Supplementary Figure 12**. (**A**) Plot of multi-carbapenemase Enterobacterales genomes from NCBI GenBank with fewer than 40 contigs ordered by number of contigs (n=2033). Width of lines corresponds to the number of genomes. A horizontal line was added at 20 contigs to indicate an inflection point. (**B**) Distribution of genomes by number of contigs. Assemblies with a long-read sequencing platform (Oxford Nanopore Technologies, PacBio) listed in their metadata are coloured dark grey, and those without are coloured light grey. A horizontal line was added at 20 contigs to indicate an inflection point.
